# Using mathematical constraints to explain narrow ranges for allele-sharing dissimilarities

**DOI:** 10.1101/2024.11.19.624404

**Authors:** Xiran Liu, Zarif Ahsan, Noah A. Rosenberg

## Abstract

Allele-sharing dissimilarity (ASD) statistics are measures of genetic differentiation for pairs of individuals or populations. Given the allele-frequency distributions of two populations—possibly the same population—the expected value of an ASD statistic is computed by evaluating the expectation of the pairwise dissimilarity between two individuals drawn at random, each from its associated allele-frequency distribution. For each of two ASD statistics, which we term 𝒟_1_ and 𝒟_2_, we investigate the extent to which the expected ASD is constrained by allele frequencies in the two populations; in other words, how is the magnitude of the measure bounded as a function of the frequency of the most frequent allelic type? We first consider dissimilarity of a population with itself, obtaining bounds on expected ASD in terms of the frequency of the most frequent allelic type in the population. We then examine pairs of populations that might or might not possess the same most frequent allelic type. Across the unit interval for the frequency of the most frequent allelic type, the expected allele-sharing dissimilarity has a range that is more restricted than the r0, 1s interval. The mathematical constraints on expected ASD assist in explaining a pattern observed empirically in human populations, namely that when averaging across loci, allele-sharing dissimilarities between pairs of individuals often tend to vary only within a relatively narrow range.

## 1. Introduction

Statistics based on concepts of allele-sharing dissimilarity (ASD) [Mountain and Cavalli-Sforza, 1997, Mountain and Ramakrishnan, 2005, Gao and Martin, 2009] are important tools in population-genetic data analysis. Beginning with the alleles of two diploid individuals at a genetic locus, a function of the four alleles is computed, producing a value ranging from 0 for the minimum dissimilarity to 1 for the maximum. Among genetic dissimilarity measures, ASD-based statistics are relatively easy to describe and compute. They are meaningful for pairs of individuals, or—if many individuals are considered—pairs of populations, or an individual and a population. Hence, features of allele-sharing dissimilarities are often used for understanding genetic variation within and among populations [Mountain and Cavalli-Sforza, 1997, Mountain and Ramakrishnan, 2005, Witherspoon et al., 2007, Rosenberg, 2011].

Studies of population-genetic statistics that consider dissimilarities across individuals suggest that ranges of observed numerical values of dissimilarity statistics—notably those based on the classic statistic *F*_*ST*_ —depend in predictable ways on allele-frequency distributions [Jakobsson et al., 2013, Edge and Rosenberg, 2014, Alcala and Rosenberg, 2017, 2019, 2022]. Consider two populations, each with an allele-frequency distribution at a locus, and consider a bounded statistic that measures the dissimilarity of the two populations as a function of the allele frequencies. Although the statistic is bounded, typically in r0, 1s, tighter constraints might exist on the dissimilarity in terms of the separate frequency distributions in the two populations. A value such as 0.55 or 0.7 might then be appropriate to interpret not in relation to the entire unit interval, but in relation to a shorter interval suited to its allele frequencies. Such interpretations have been used to explain unexpected numerical patterns in *F*_*ST*_ —such as a low *F*_*ST*_ value among high-diversity African populations [Jakobsson et al., 2013], and a high *F*_*ST*_ among chimpanzee populations relative to its value between chimpanzees and humans [Alcala and Rosenberg, 2022].

Empirical findings suggest that allele-sharing dissimilarities are also constrained by allele frequencies. For example, the values of allele-sharing dissimilarities have been seen to be quite similar across many computations. Consider, for example, the computations of allele-sharing dissimilarities as averages across many loci for pairs of individuals in Figure 5A of Rosenberg [2011]. In those computations, which consider genome-wide loci in diverse human populations, most pairs of individuals possess allele-sharing dissimilarities between 0.55 and 0.7. Does the narrow range arise from mathematical constraints on ASD measures in relation to the allele frequencies?

We have recently investigated the mathematical properties of two formulations of population-level ASD measures, exploring mathematical properties of the expected genetic dissimilarity between pairs of individuals sampled within and between populations [Liu et al., 2023]. Here, building upon our previous mathematical results, we derive bounds on the two expectations, both within and between populations, in two scenarios: first, when the number of allelic types at a given locus is fixed at *I* and the allele-frequency distributions within a population can be arbitrary among these *I* alleles; second, when both *I* and the frequency *M* of the most frequent allelic type within a population are held constant. In both scenarios, we focus on the upper and lower bounds on the genetic dissimilarities in terms of the frequency of the most frequent allelic type. We find that indeed, ASD values are mathematically constrained to subintervals of r0, 1s, and that the constraints can assist in explaining features of ASD in human populations.

## 2 Preliminaries

### 2.1 Definitions

Following Liu et al. [2023], we consider two variants of the ASD concept, which we denote by 𝒟_1_ and 𝒟_2_. For 𝒟_1_, “allele-sharing” for two diploid individuals is interpreted as the number of shared elements in their sets of alleles. 𝒟_1_ then uses 1 minus the normalized count of the shared alleles as the dissimilarity. Consider a locus with four distinct alleles, A, B, C, and D, the minimum number required so that all possible cases for diploid genotypes exist. Two individuals both with genotype AB have 2 alleles shared, and 𝒟_1_ = 0. An individual with genotype AB and an individual with genotype AC have 1 shared allele, namely A, and 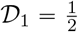.

𝒟_2_ instead considers alleles individually, evaluating the fraction of pairs of alleles, one from the first individual and one from the second, that are distinct. For two individuals with genotype AB, 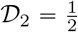 among the four possible pairs of alleles — (A,A), (A,B), (B,A), and (B,B), where the first entry in the pair represents an allele from the first individual and the second entry is an allele from the second — two of four contain distinct alleles. An individual with genotype AB and an individual with genotype AC have 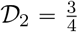.

Consider a locus with *I* ⩾ 2 allelic types, and suppose the allele frequencies in a population are **p** = (*p*_1_, *p*_2_, …, *p*_*I*_), where *p*_*i*_ represents the frequency of allele *i*. The frequencies satisfy 0 ⩽ *p*_*i*_ ⩽ 1 for all *i*, and 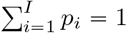. Without loss of generality, let *p*_1_ = *M* represent the largest entry in the allele-frequency vector (*p*_1_, *p*_2_, …, *p*_*I*_). When we consider allele-frequency vectors in two populations, we let population 2 have allele frequencies **q** = (*q*_1_, *q*_2_, …, *q*_*I*_) satisfying 0 ⩽ *q*_*i*_ ⩽ 1 for all *i*, and 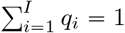. We define

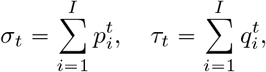

for *t* =1, 2, 3, 4, where *σ*_1_ = *τ*_1_ = 1. We also define

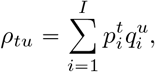

where (*t, u*) is equal to (1, 1), (1, 2), (2, 1), or (2, 2).

We denote the dissimilarity 𝒟 between two individuals within the same population with allele-frequency vector **p** by 𝒟^*w*^(**p**); here, 𝒟 is understood to refer to one of the two dissimilarities, 𝒟_1_ or 𝒟_2_. We denote the corresponding dissimilarity between two individuals from different populations with allele-frequency vectors **p** and **q** by 𝒟^*b*^(**p, q**). We often drop the arguments for convenience.

### 2.2 Review of ASD mathematical results

In Liu et al. [2023], we studied a probabilistic model in which individuals are randomly sampled from allele-frequency distributions and 𝒟_1_ and 𝒟_2_ are computed. The expected value of 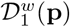 satisfies
*

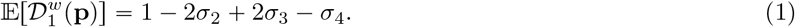

For *I* =2, substituting *p*_2_ = 1 − *p*_1_ so that 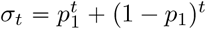, Eq. 1 becomes 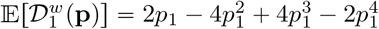. We also have

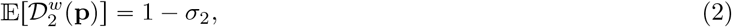

and for the *I* = 2 case, Eq. 2 simplifies to 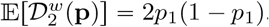.

For the between-population dissimilarity 𝒟^*b*^(**p**), we obtain

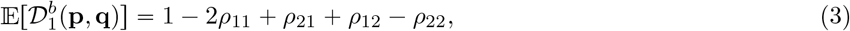

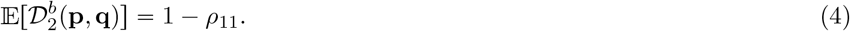

Eqs. 1, 2, 3, and 4 correspond to Eqs. 3, 9, 16, and 22 of Liu et al. [2023].

### 2.3 Review of majorization theory

We recall some results from majorization theory that will assist in finding bounds on ASD statistics. Majorization describes partial orderings on vectors with a shared sum.

#### Definition 2.1

(Majorization, 1.A.1 of Marshall et al. [2010]). Vector **x** ∈ ℝ^*n*^ is said to *majorize* vector **y** ∈ ℝ^*n*^ if, when the components of **x** and **y** are each rearranged in non-increasing order, (i) 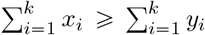 for all *k* = 1, 2, … *n* −1 and (ii) 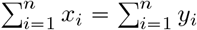. Equivalently, **y** is said to be *majorized* by **x**.

If **x** majorizes **y**, then we write **x** ≻ **y**. Functions that preserve majorization order are said to be Schur-convex.

#### Definition 2.2

(Schur-convexity, 3.A.1 of Marshall et al. [2010]). A function *f* : ℝ^*n*^ →ℝ is said to be *Schur-convex* if **x** ≻ **y** implies *f* (**x**) ⩾ *f* (**y**). The function is *strictly Schur-convex* if **x** ≻ **y** and **x** ≠ **y** implies *f* (**x**) > *f* (**y**). A function *f* is *Schur-concave* if −*f* is Schur-convex and *strictly Schur-concave* if −*f* is strictly Schur-concave.

#### Theorem 2.3

(Schur convexity condition, 3.A.4 of Marshall et al. [2010]). *Let* ℐ ⊂ ℝ *be an open interval and let f* : ℐ^*n*^ → ℝ *be a continuously differentiable function. Function f is Schur-convex if and only if f is symmetric in its n arguments and for all* (*i, j*) *with* 1 ⩽ *i, j* ⩽ *n*,

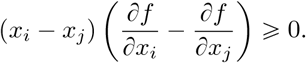

*Further, if equality requires x*_*i*_ = *x*_*j*_, *then f is strictly Schur-convex*.

*Similarly, f is Schur-concave if and only if f is symmetric in its n arguments and for all* (*i, j*) *with* 1 ⩽ *i, j* ⩽ *n*,

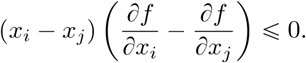

*If equality requires x*_*i*_ = *x*_*j*_, *then f is strictly Schur-concave*.

Denote the unit (*I* − 1)-simplex by Δ^*I*−1^:

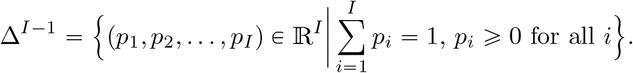

#### Proposition 2.4

(Majorization inequality for a unit simplex, Section 2.2 of Aw and Rosenberg [2018]). *For all vectors* **p** *in the unit* (*I* − 1)*-simplex* ΔΔ^*I*−1^,

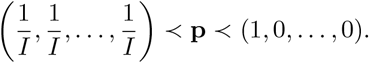

#### Proposition 2.5

(Majorization inequality for vectors in the simplex, with a specified value of the largest entry; see the proof of Theorem 3.9 of Aw and Rosenberg [2018]). *Let* **p** *be a vector of length I chosen within the simplex, with largest largest entry equal to M*; *that is*, **p** ∈ Δ^*Ĩ*−1^, *where*

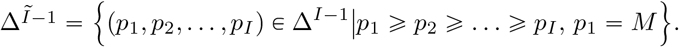

*Then*

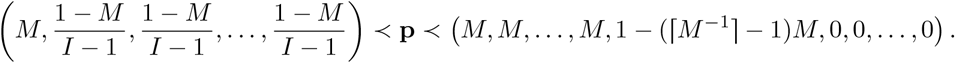

Here, the left-hand vector has *I* − 1 entries equal to 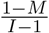. The right-hand vector has ⌈*M* ^−1^⌉− 1 entries equal to *M* followed by an entry of 1 − (⌈*M* ^−1^⌉− 1)*M* and zeroes for the remaining entries. For convenience, we write **p**_min_ = (*M*, …, *M*, 1 − (⌈*M* ^−1^⌉− 1)*M*, 0, …, 0) and 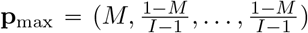,), noting that because we will be working with Schur-concave functions, the most majorized vector becomes the “maximum.”

#### Theorem 2.6

(Rearrangement inequality, Theorem 368 of Hardy et al. [1952]). *Consider two sets of I real numbers a*_1_ ⩾ *a*_2_ ⩾ … ⩾ *a*_*I*_ *and b*_1_ ⩾ *b*_2_ ⩾ … ⩾ *b*_*I*_. *For each permutation b*_*σ*(1)_, *b*_*σ*(2)_, …, *b*_*σ*(*I*)_ *of b*_1_, *b*_2_, …, *b*_*I*_,

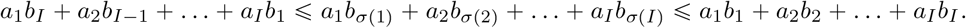

## 3 Mathematical constraints on within-population dissimilarity

Using Eqs. 1 and 2, we consider two sets of mathematical constraints on the within-population dissimilarity measures 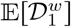 and 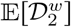. First, fixing the number of allelic types *I* but permitting the allele-frequency distribution to be arbitrary, we consider general bounds as functions of *I*. Second, because the largest allele frequency *M* might impose further restrictions on the allele-frequency distribution, we consider the bounds when fixing both *I* and *M*.

### 3.1 Bounds on 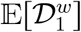 when the number of allelic types *I* is*I* fixed

Let 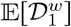 be a function of **p** = (*p*_1_, *p*_2_, …, *p*_*I*_) following Eq. 1, where **p** ∈ Δ^−1^, the standard (*I* − 1)-simplex. Denote 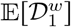 by *f* (**p**).

#### Lemma 3.1.

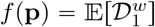, *as a function of* **p**∈ Δ^−1^*is strictly Schur-concave*.

The proof of the lemma appears in Appendix A. Using the strict Schur-concavity of the function 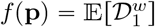 from Lemma 3.1, we arrive at the following theorem.

#### Theorem 3.2.

*Suppose without loss of generality that p*_1_ ⩾ *p*_2_ ⩾ … ⩾ *p*_*I*_ *for* **p** = (*p*_1_, *p*_2_, …, *p*_*I*_). *Then*

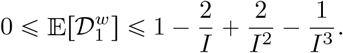

*Equality at the lower bound is reached if and only if p*_1_ = 1 *and p*_*i*_ = 0 *for* 2 ⩽ *i* ⩽ *I. Equality at the upper bound is reached if and only if* 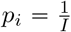 *for* 1 ⩽ *i* ⩽ *I*.

*Proof*. By Proposition 2.4, 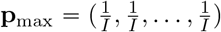 is majorized by all and **p**_min_ = (1, 0, …, 0) majorizes all **p**∈ Δ^−1,^ Because 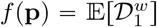 is strictly Schur-concave by Lemma 3.1, by definition of strict Schur-concavity (Definition 2.2), *f* (**p**_max_) ⩾ *f* (**p**) for all **p**∈ Δ^−1^and *f* (**p**_min_) ⩽ *f* (**p**) for all **p**∈ Δ^−1.^Therefore,

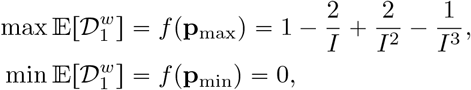

with equality if and only if **p** lies at the specified points.

In the simplest case of *I* = 2, we find that 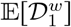 is maximized for 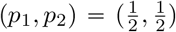, with 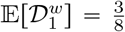. It is minimized for (*p*_1_, *p*_2_) = (1, 0), at which 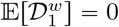.

The relationship between the upper bound on 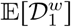 and *I* is shown in Figure 1. The figure shows a strictly increasing sequence, as is clear by noting that the derivative of the upper bound of 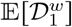 as a function of *I* is (2*I*^2^ − 4*I* + 3)/*I*^4^, a strictly positive function for *I* ⩾ 2. As *I* → ∞, this upper bound approaches 1.

**Figure 1:**
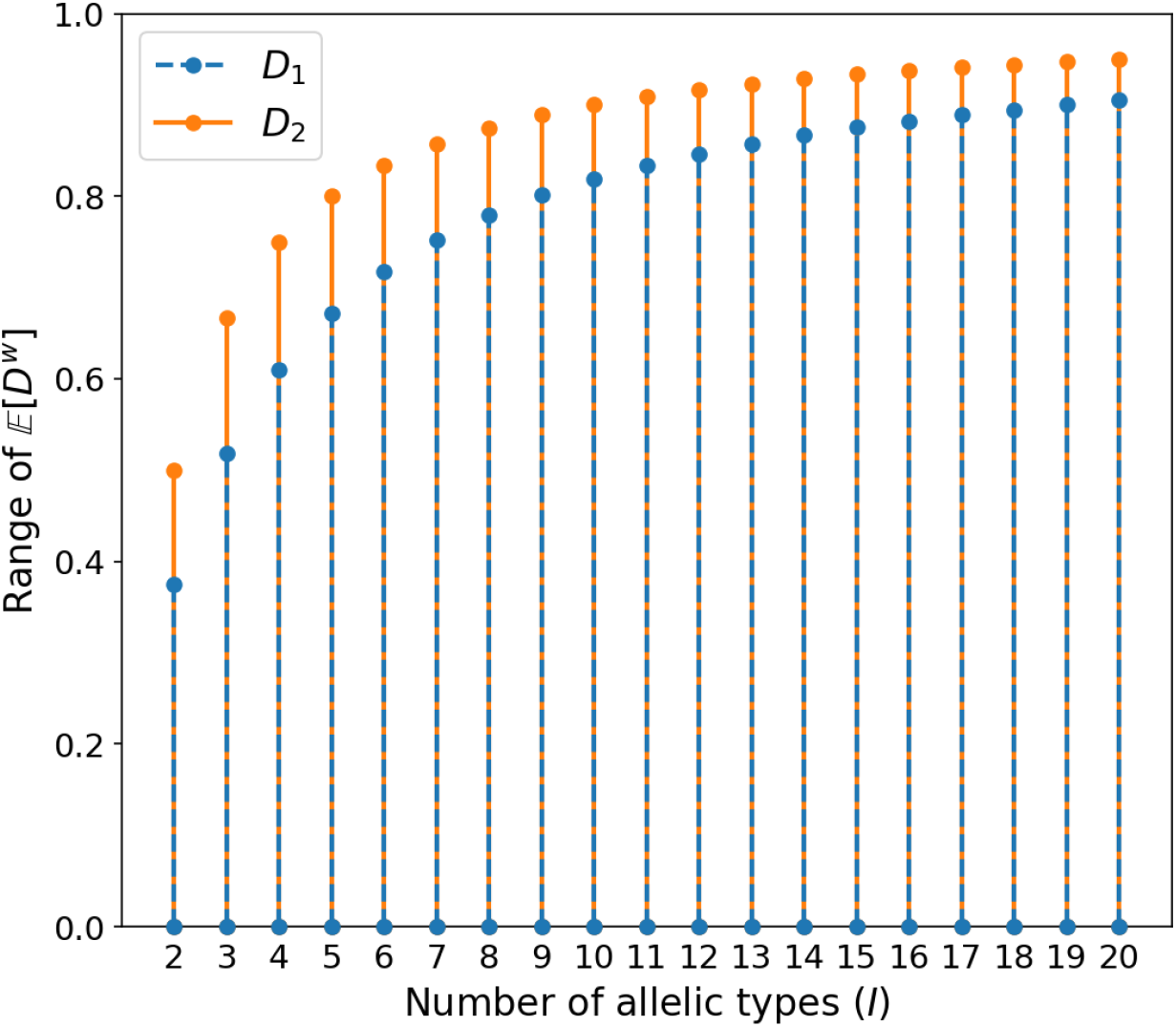
Range of 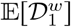 and 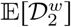 as functions of the number of allelic types *I* when the allele-frequency vector is permitted to be arbitrary, as stated in Theorems 3.2 and 3.6.

### 3.2 Bounds on 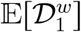 when the largest allele frequency is fixed

If the largest allele frequency *M* is fixed, then a tighter constraint is imposed on the range of values that 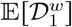 can take. To derive this constraint, we use Proposition 2.5.

#### Theorem 3.3.

*Suppose p*_1_ ⩾ *p*_2_ ⩾ … ⩾ *p*_*I*_, *and suppose p*_1_ = *M is fixed*, 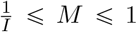 *Let* 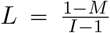 *and* 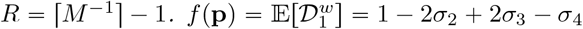 *σ*_4_ *is bounded by*

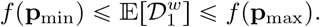

*Equality with the lower bound is achieved if and only if* **p** = **p**_min_ = (*M*, …, *M*, 1−(⌈*M* ^−1^⌉−1)*M*, 0, …, 0), *producing f* (**p**_min_) = 1−*R*(2*M* ^2^−2*M* ^3^+*M* ^4^)−2(1−*RM*)^2^+2(1−*RM*)^3^−(1−*RM*)^4^. *Equality with the upper bound is achieved if and only if* 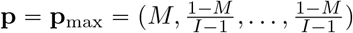, *producing f* (**p**_max_) = 1−2*M* ^2^+2*M* ^3^ −*M* ^4^ −(*I* −1)(2*L*^2^ −2*L*^3^ +*L*^4^).

The proof is straightforward: by Proposition 2.5, **p**_min_ ≻ **p** ≻ **p**_max_ for all **p**∈ Δ^*Ĩ*−1^. Because *f* is strictly Schur-concave (Lemma 3.1), by the definition of Schur-concavity (Definition 2.2), *f* (**p**_min_) ⩽ *f* (**p**) ⩽ *f* (**p**_max_) for all **p**∈ Δ^*Ĩ*−1^, with the appropriate equality conditions.

In the *I* = 2 case, there is a single choice for **p**, and the two bounds coincide. For each *I* from 2 to 9, Figure 2 plots the region specified by the theorem, illustrating that as *I* increases, the size of the permissible region grows.

**Figure 2:**
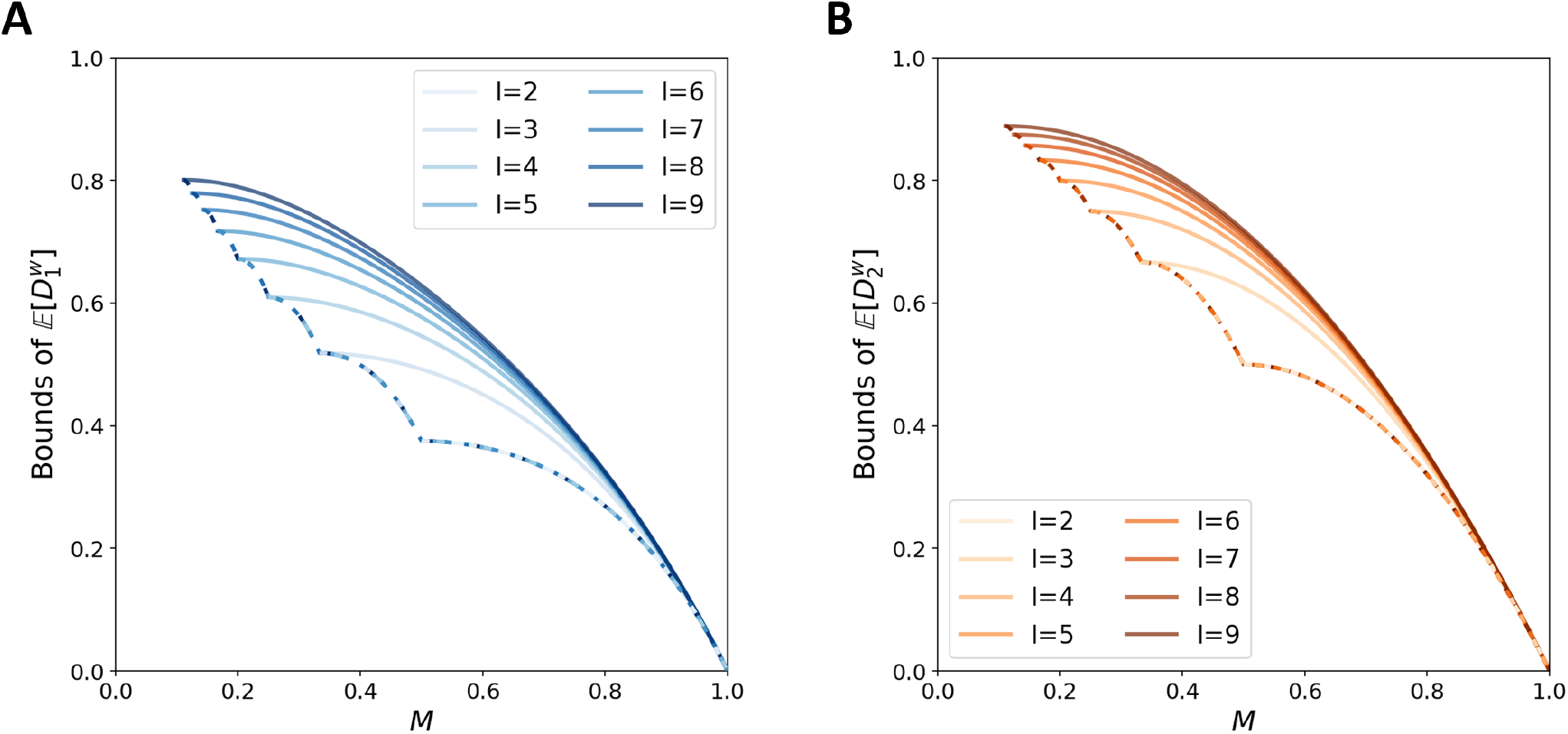
Bounds of expected dissimilarities for *I* = 2 to 9 allelic types when the largest allele frequency is fixed to be *M*, as stated in Theorems 3.3 and 3.7. (A) 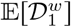 (B) 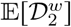 The solid line corresponds to the upper bound, and the dashed line corresponds to the lower bound, with lower bounds for different *I* values overlapping.

The vector **p** that produces equality of the lower bound of 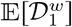 given *M* is exactly the same as the one that minimizes the heterozygosity given *I* and *M*; similarly, the vector that produces equality of the upper bound of 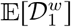 given *M* is the vector tht maximizes heterozygosity given *I* and *M* [Reddy and Rosenberg, 2012].

#### Proposition 3.4.

*With fixed I, the region bounded by the upper and lower bounds on* 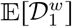 *as a function of M has area*

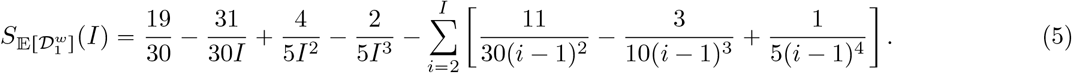

The proof appears in Appendix B. Letting *I* → ∞ in Eq. 5, noting that the Riemann zeta function satisfies 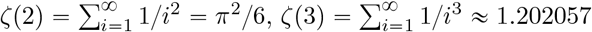, and 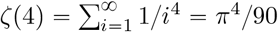, the area approaches

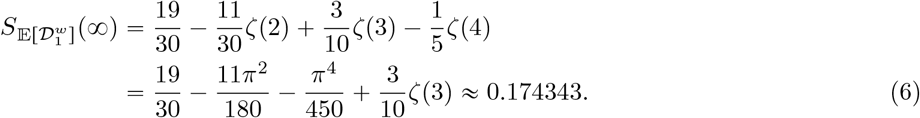

The area of the region is plotted as a function of *I* in Figure 3.

**Figure 3:**
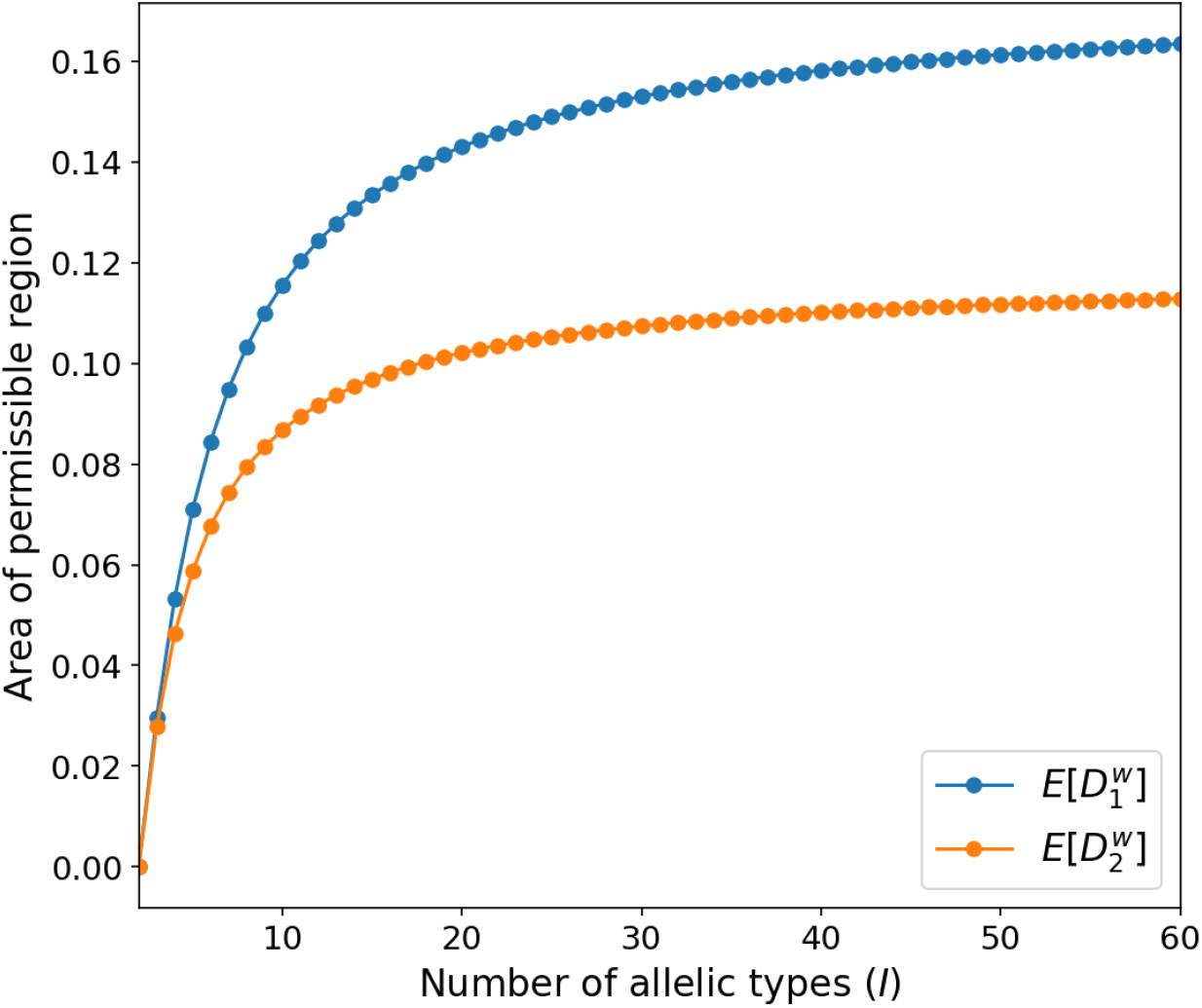
Area of the permissible region for expected dissimilarities for *I* = 2 to 60 when the largest allele frequency is fixed to be *M* and *M* ranges from 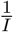 to 1, as stated in Propositions 3.4 and 3.8.

### 3.3 Bounds on 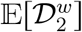 when the number of allelic types *I* is fixed

Bounds on 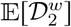 can be obtained similarly to those on 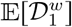. We write 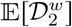 as a function *g*(**p**) following Eq. 2, where **p**Δ^*I*−1^ The functional form 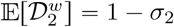 admits known results for the homozygosity *σ*_2_.

#### Lemma 3.5.

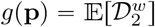, *as a function of* **p**Δ^*I*−1^, *is strictly Schur-concave*.

The proof of the lemma follows directly from the strict Schur-convexity of homozygosity *σ*_2_ [Aw and Rosenberg, 2018, p. 720]. As a function of **p**, *g*(**p**) = 1 − *σ*_2_, so that *g*(**p**) is strictly Schur-concave by Definition 2.2.

#### Theorem 3.6.

*Suppose without loss of generality that p*_1_ ⩾ *p*_2_ ⩾ … ⩾ *p*_*I*_ *for* **p** = (*p*_1_, *p*_2_, …, *p*_*I*_). *Then*

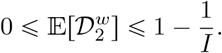

*Equality at the lower bound is reached if and only if p*_1_ = 1 *and p*_*i*_ = 0 *for* 2 ⩽ *i* ⩽ *I. Equality at the upper bound is reached if and only if* 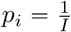 *for* 1 ⩽ *i* ⩽ *I*.

*Proof*. Using the strict Schur-concavity of function *g*, the proof follows that of Theorem 3.2. We obtain

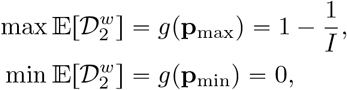

with equality if and only if **p** lies at the specified points.

In the *I* = 2 case, we have a maximum value of 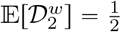 if and only if 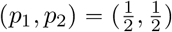 and a minimum value of 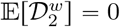 if and only if (*p*_1_, *p*_2_) = (1, 0).

The relationship between the upper bound on 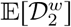 and *I* appears in Figure 1. It is a strictly increasing sequence, and as *I* → ∞, the upper bound approaches 1. Note that for *I* ⩾ 2, the upper bound on 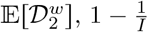, strictly exceeds the upper bound on 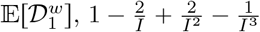 as 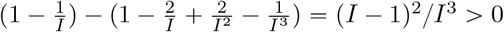.

### 3.4 Bounds on 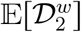 when the largest allele frequency is fixed

The bounds on 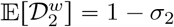 when *p*_1_ = *M* restate known bounds on *σ*_2_.

#### Theorem 3.7.

*Suppose p*_1_ ⩾ *p*_2_ ⩾ … ⩾ *p*_*I*_, *and suppose p*_1_ = *M is fixed*, 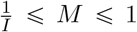 *Let* 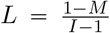 *and* 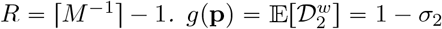 *is bounded by*

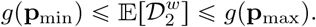

*Equality with the lower bound is achieved if and only if* **p** = **p**_min_ = (*M*, …, *M*, 1 − (⌈*M* ^−1^⌉ − 1)*M*, 0, …, 0), *producing g*(**p**_min_) = 1 − *RM* ^2^ − (1 − *RM*)^2^. *Equality with the upper bound is achieved if and only if* 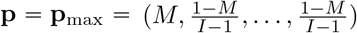, *producing g*(**p**_max_) = 1 − *M* ^2^ −(*I* − 1)*L*^2^.

The theorem is a restatement of Theorem 2 in Reddy and Rosenberg [2012], which provided the bounds on *σ*_2_ for fixed *I* and *M*. In the *I* = 2 case, the upper and lower bounds coincide. The upper and lower bounds are achieved at precisely the same allele-frequency vectors that achieve the upper and lower bounds on 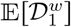.

The relationships between the lower and upper bounds of 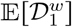 and 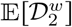 and the frequency *M* appear in Figure 2. Both bounds of 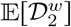 exceed than those of 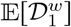, as is clear by noting that 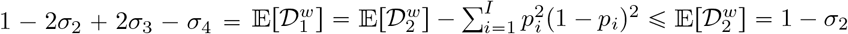.

#### Proposition 3.8.

*With fixed I, the region bounded by the upper and lower bounds of* 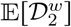 *as a function of M has area*

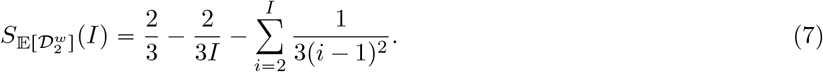

As *I* → ∞, the area approaches

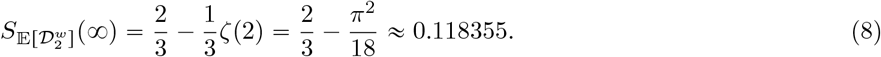

Proposition 3.8 and the calculation of 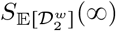 restate Proposition 16 of Reddy and Rosenberg [2012] for the bounds on *σ*_2_. The area of the region is plotted as a function of *I* in Figure 3.

## 4 Mathematical constraints on between-population dissimilarity

Next, we look at the bounds for the between-population dissimilarities, which involve the allele-frequency vectors of two populations, **p** and **q**.

### 4.1 Bounds on 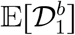 when the number of allelic types is fixed

If the number of distinct alleles is fixed and the allele-frequency distributions can be arbitrary, then the bounds on both between-population dissimilarity measures are trivial.

#### Proposition 4.1.

*Suppose without loss of generality that p*_1_ ⩾ *p*_2_ ⩾ … ⩾ *p*_*I*_ *for* **p** = (*p*_1_, *p*_2_, …, *p*_*I*_), *and no constraints are placed on* **q**. *Then*

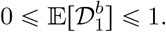

*Equality at the lower bound is reached if and only if p*_1_ = *q*_1_ = 1 *and p*_*i*_ = *q*_*i*_ = 0 *for* 2 ⩽ *i* ⩽ *I. Equality at the upper bound is reached if and only if* **p** *and* **q** *satisfy p*_*i*_*q*_*i*_ = 0 *for* 1 ⩽ *i* ⩽ *I*.

*Proof*. For a pair of individuals, one from population 1 and one from population 2, 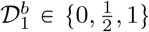,. Hence, as a function of allele-frequency vectors, we know 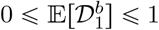.

We consider the equality condition 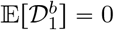. For a pair of individuals, 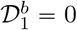 if and only if both individuals have exactly the same diploid genotype. 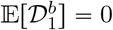 requires that all pairs of individuals, one from one population and one from the other, possess the same diploid genotype. If either population possesses at least two distinct alleles with nonzero frequency, then the probability is positive that 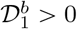 for a pair of random individuals, one from one population and one from the other, so that 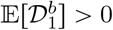. We conclude that if 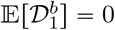, then two populations must have the same single allele. With *p*_1_ ⩾ *p*_2_ ⩾ … ⩾ *p*_*I*_, it follows that *p*_1_ = *q*_1_ = 1 and *p*_*i*_ = *q*_*i*_ = 0 for 2 ⩽ *i* ⩽ *I*.

For the equality condition 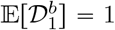, writing 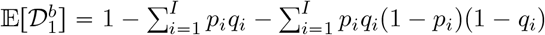, we find that 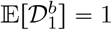 implies 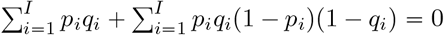, from which we conclude that the non-negative *p*_*i*_ and *q*_*i*_ bounded above by 1 must satisfy *p*_*i*_*q*_*i*_ = 0 for all *i*.

In the case of 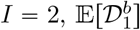 is minimized for **p** = **q** = (1, 0), and maximized for **p** = (1, 0), **q** = (0, 1).

### 4.2 Bounds on 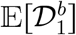 when the largest allele frequencies are fixed

We next consider scenarios in which the largest allele frequency is fixed in both populations. We first investigate the bounds on 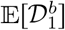 in the case that the same allelic type has the largest frequency in both populations.

#### Theorem 4.2.

*Suppose p*_1_ = max{*p*_1_, *p*_2_, …, *p*_*I*_} *and q*_1_ = max{*q*_1_, *q*_2_, …, *q*_*I*_}. *Suppose without loss of generality that p*_1_ ⩾ *p*_2_ ⩾ … ⩾ *p*_*I*_. (*i*) *If p*_1_ = *M*_1_ *and q*_1_ = *M*_2_ *are fixed, then*

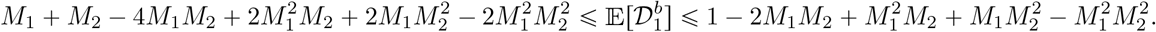

(*ii*) *Equality with the upper bound is achieved if and only if for all i*, 2 ⩽ *i* ⩽ *I, p*_*i*_ = 0 *or q*_*i*_ = 0. (*iii*) *Equality with the lower bound is achieved if and only if* 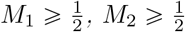, *and* 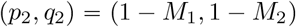.

The proof appears in Appendix C. Note that if 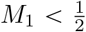 or 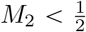, we have not obtained strict inequalities; the bounds in the theorem hold, but the lower bound is not the strictest possible inequality. The bounds in the theorem are depicted in Figure 4A-F.

**Figure 4:**
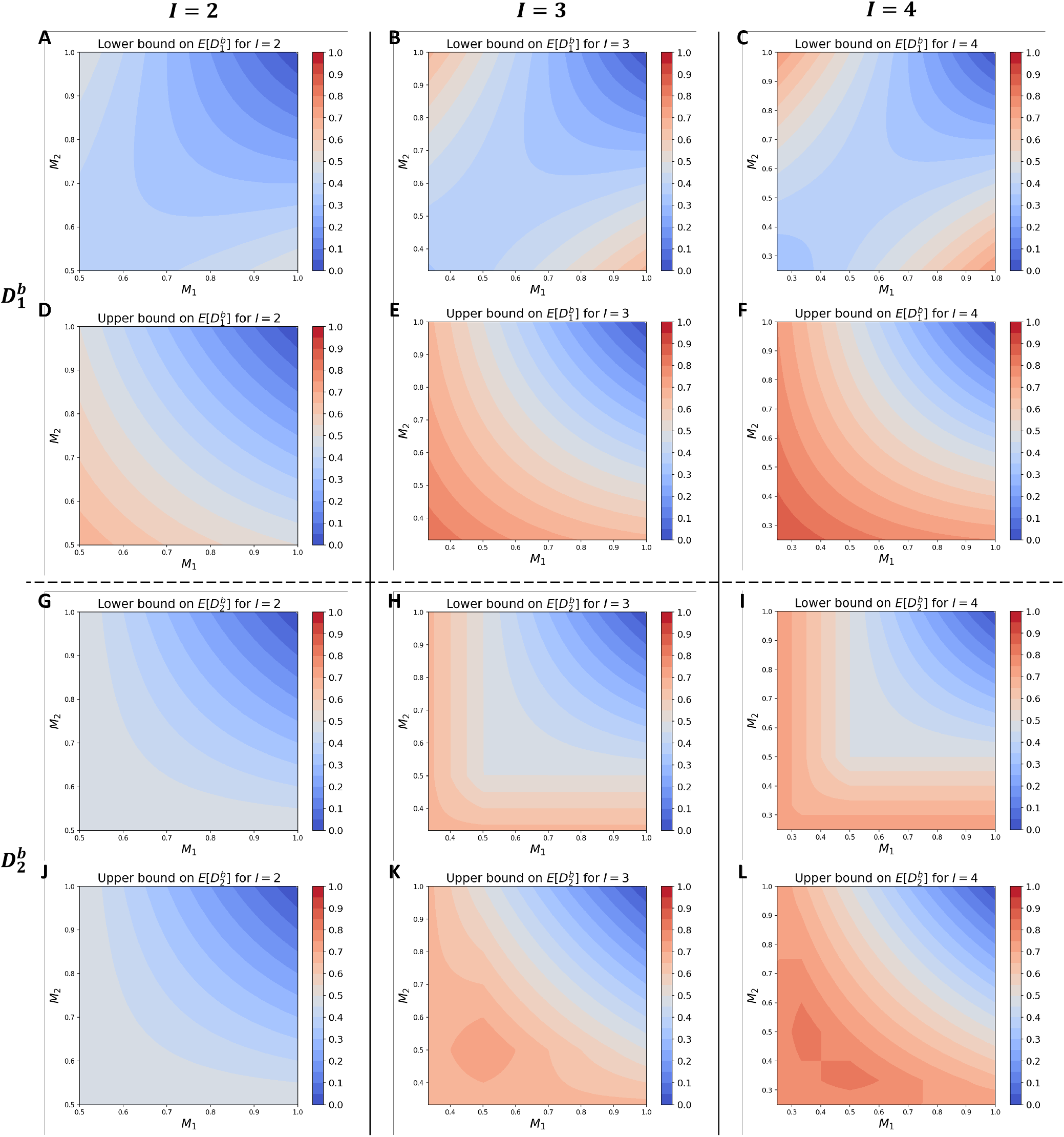
Bounds for 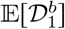 and 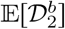 for *I* = 2 to 4 when two populations share the most frequent allelic type. The frequencies of the most frequent allelic type are fixed to be *M*_1_ and *M*_2_ in the two populations. (A) 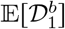, lower bound for *I* = 2. (B) 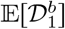, lower bound for *I* = 3. (C) 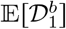, lower bound for *I* = 4. (D) 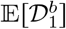, upper bound for *I* = 2. (E) 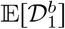, upper bound for *I* = 3. (F) 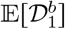, upper bound for *I* = 4. (G) 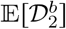, lower bound for *I* = 2. (H) 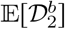, lower bound for *I* = 3. (I) 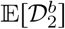, lower bound for *I* = 4. (J) 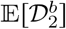, upper bound for *I* = 2. (K) 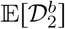, upper bound for *I* = 3. (L) 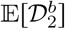, upper bound for *I* = 4. Bounds for 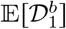 follow Theorem 4.2. Bounds for 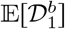 follow Corollary 4.6. The lower bound for 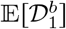 is loose if 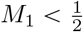 or 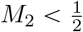. The lower bound for 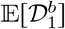 if 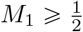 and 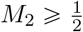, the upper bound for 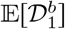, and the bounds for 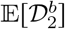 are strict.

For a general scenario in which two populations might have different allelic types for their most frequent allele, we also obtain loose inequalities. We proceed by first bounding 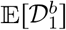 in relation to 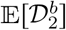, then deriving bounds for 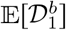 based on the bounds we obtain for 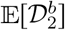 in Section 4.4.

#### Theorem 4.3.

*Suppose* max{*p*_1_, *p*_2_, …, *p*_*I*_} = *M*_1_ *and* max{*q*_1_, *q*_2_, …, *q*_*I*_} = *M*_2_. *Suppose without loss of generality that p*_1_ ⩾ *p*_2_ ⩾ … ⩾ *p*_*I*_. *Then*

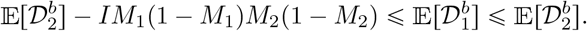

The proof appears in Appendix D. The bounds in the theorem are depicted in Figure 5A-F. Together with the lower and upper bounds for 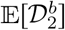 in Section 4.4, we are able to obtain a lower bound and upper bound for 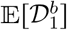.

**Figure 5:**
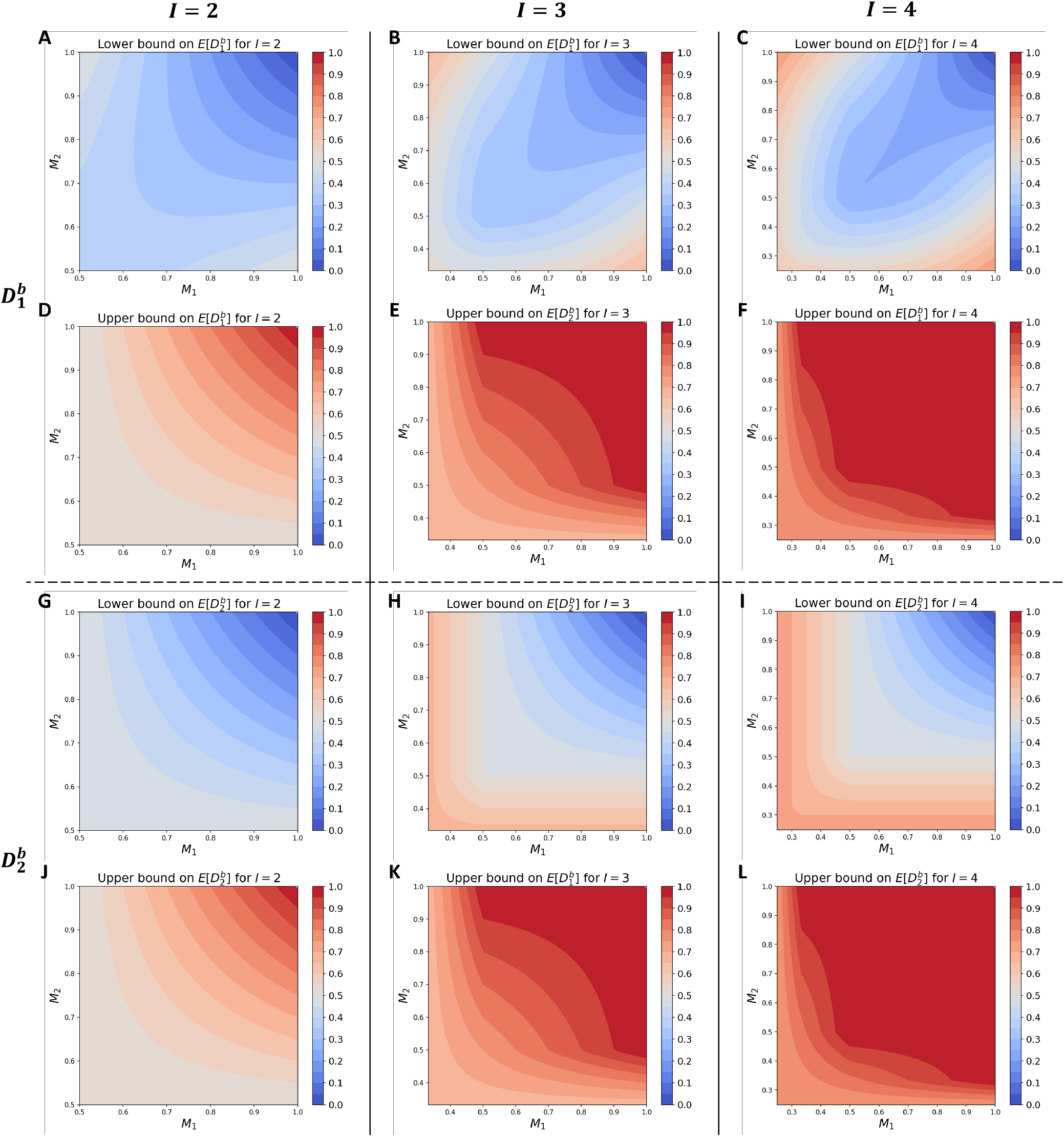
Bounds for 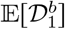 and 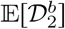 for *I* = 2 to 4 when two populations do not necessarily share the most frequent allelic type. The frequencies of the most frequent allelic type are fixed to be *M*_1_ and *M*_2_ in the two populations. (A) 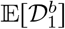, lower bound for *I* = 2. (B) 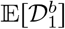, lower bound for *I* = 3. (C) 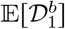, lower bound for *I* = 4. (D) 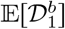, upper bound for *I* = 2. (E) 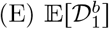, upper bound for *I* = 3. (F) 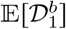, upper bound for *I* = 4. (G) 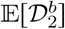, lower bound for *I* = 2. (H) 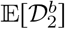, lower bound for *I* = 3. (I) 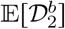, lower bound for *I* = 4. (J) 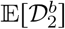, upper bound for *I* = 2. (K) 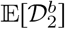, upper bound for *I* = 3. (L) 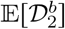, upper bound for *I* = 4. Bounds for 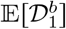 follow Theorem 4.3. Bounds for 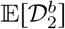 follow Theorem 4.5. Bounds for 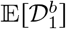 are loose and bounds for 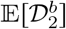 are strict.

### 4.3 Bounds on 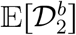 when the number of allelic types is fixed

As we observed with 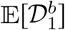, the bounds on 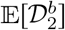 are trivial when the number of distinct alleles is fixed a nd the allele-frequency distribution can be arbitrary.

#### Proposition 4.4.

*Suppose without loss of generality that p*_1_ ⩾ *p*_2_ ⩾ … ⩾ *p*_*I*_ *for* **p** = (*p*_1_, *p*_2_, …, *p*_*I*_), *and no constraints are placed on* **q**. *Then*

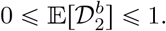

*Equality at the lower bound is reached if and only if p*_1_ = *q*_1_ = 1 *and p*_*i*_ = *q*_*i*_ = 0 *for* 2 ⩽ *i* ⩽ *I. Equality at the upper bound is reached if and only if* **p** *and* **q** *satisfy p*_*i*_*q*_*i*_ = 0 *for* 1 ⩽ *i* ⩽ *I*.

*Proof*. We have 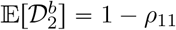, and 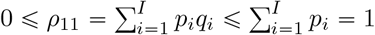, so that 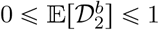.

The equality condition 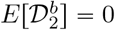 holds if and only if *ρ*_11_ = 1; *ρ*_11_ = 1 implies 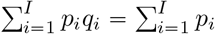, from which 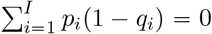 and *q*_*i*_ = 1 for each *i* for which *p*_*i*_ > 0. Symmetrically, 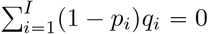 and *p*_*i*_ = 1 for each *i* for which *q*_*i*_ > 0. We conclude that for all *i*, (*p*_*i*_, *q*_*i*_) = (1, 1) or (0, 0). Because *p*_1_ ⩾ *p*_2_ ⩾ …⩾ *p*_*I*_, we have (*p*_1_, *q*_1_) = (1, 1) and (*p*_*i*_, *q*_*i*_) = (0, 0) for 2 ⩽ *i* ⩽ *I*.

The equality condition 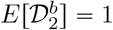 holds if and only if *ρ*_11_ = 0, so that *p*_*i*_*q*_*i*_ = 0 for all *i*.

For 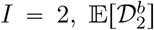 has the same minimum and maximum as 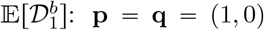 for the minimum and **p** = (1, 0), **q** = (0, 1) for the maximum.

### 4.4 Bounds on 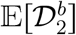 when the largest allele frequencies are fixed

Unlike for 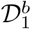, we obtain strict bounds 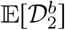 in the general scenario in which two populations may have different allelic types for the most frequent allele, with frequencies *M*_1_ and *M*_2_ respectively. We recall the rearrangement inequality (Theorem 2.6) and use a related Lemma E.3 in Appendix E.

#### Theorem 4.5.

*Suppose* max{*p*_1_, *p*_2_, …, *p*_*I*_} = *M*_1_ *and* max{*q*_1_, *q*_2_, …, *q*_*I*_} = *M*_2_. *Suppose without loss of generality that p*_1_ ⩾ *p*_2_ ⩾ … ⩾ *p*_*I*_.

i. 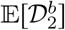 *as a function of* **p** *and* **q**, *denoted by ℓ*(**p, q**), *is bounded by*

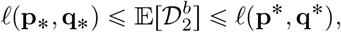

*for particular vectors* **p**_∗_, **q**_∗_, **p**\*, *and* **q**\*.
ii. *Equality at the lower bound is reached if*

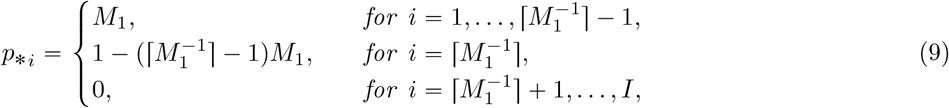

*and*

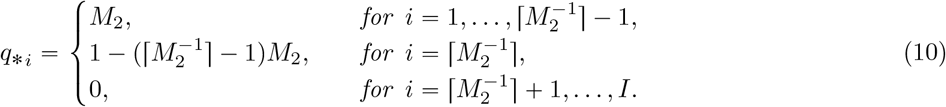

*The minimum value is*

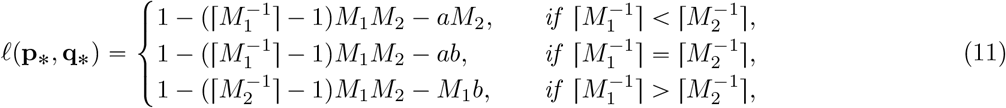

*where* 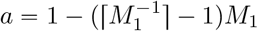 *and* 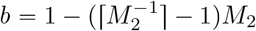.
iii. *Equality at the upper bound is reached if* **p**\* = **p**_∗_ (*Eq. 9*), *and*

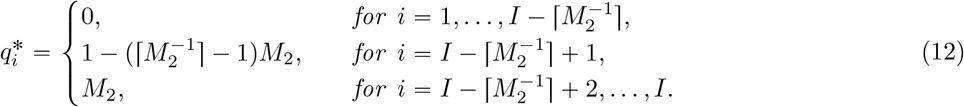

*The maximum value is*

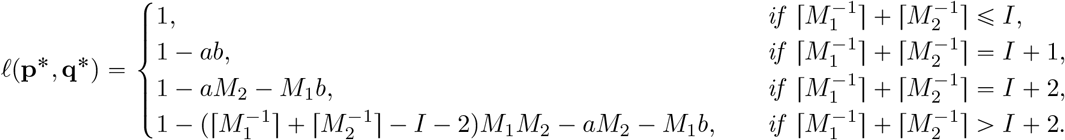

The proof of the theorem appears in Appendix E. The bounds in the theorem are depicted in Figure 5G-L. The bounds also appear in the loose bounds for 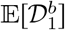 in Theorem 4.3 in the same setting, with 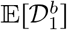 bounded above by the upper bound on 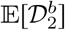 and below by the lower bound on 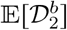 minus some additional terms.

As a corollary, we obtain the bounds for the specific scenario in which the same allelic type is most frequent in the two populations.

#### Corollary 4.6.

*Suppose p*_1_ = max{*p*_1_, *p*_2_, …, *p*_*I*_} = *M*_1_ *and q*_1_ = max{*q*_1_, *q*_2_, …, *q*_*I*_} = *M*_2_. *Suppose without loss of generality that p*_1_ ⩾ *p*_2_ ⩾ … ⩾ *p*_*I*_.

i. 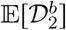 *as a function of* **p** *and* **q**, *denoted by ℓ*(**p, q**), *is bounded by*

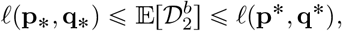

*for particular vectors* **p**_∗_, **q**_∗_, **p**\*, *and* **q**\*.
ii. *Equality at the lower bound is reached if* **p**_∗_ *follows Eq. 9 and* **q**_∗_ *follows Eq. 10. The minimum value follows Eq. 11*.
iii. *Equality at the upper bound is reached if* **p**\* = **p**_∗_ (*Eq. 9*) *and*

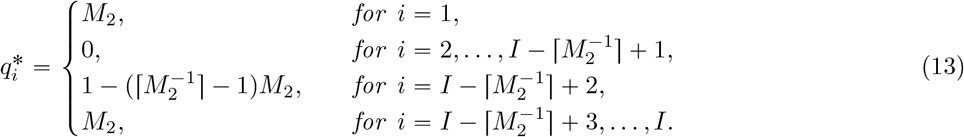

*The maximum value is*

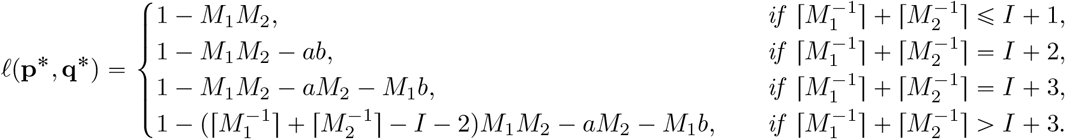

The corollary is proven in Appendix F. The bounds in the theorem are depicted in Figure 4G-L.

## 5 Data analysis

To investigate how the mathematical bounds with respect to the largest allele frequency affect the values of allele-sharing dissimilarity measures in an empirical setting, we compute the dissimilarities in a dataset of multiallelic loci in human populations.

### 5.1 Data

We analyze microsatellite genotypes in the H1048 subset of the Human Genome Diversity Project (HGDP-CEPH panel), considering 1048 individuals in 53 populations, typed at 783 microsatellite loci [Rosenberg et al., 2005, Rosenberg, 2006]. For some analyses, we restrict attention to 30 populations with sample size strictly larger than 15, considering a total of 813 individuals. For each locus, individuals with missing data are removed prior to the calculation of genetic dissimilarities for the locus. The dataset is the same as in the analysis of Liu et al. [2023].

### 5.2 Within-population dissimilarities

For each population and each locus, we compute both the theoretical and the empirical expectations 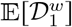 and 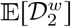. The number of population–locus combinations is 30 × 783 = 23, 490. The theoretical expectation of 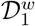 is computed by first calculating the allele frequencies of a population and then applying Eq. 1. The empirical expectation of 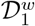 is computed by enumerating all pairs of individuals in the population, calculating their 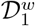 dissimilarity, and averaging over all pairs. The calculation for 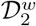 follows the same process, with Eq. 2. The calculation of theoretical and empirical expectations follows Liu et al. [2023].

We classify each locus by the number of allelic types; considering *I* from 4 to 14, the number of population–locus combinations is 630 × 30 = 18, 900, with a minimum count of 1 × 30 = 30 for *I* = 4 and a maximum count of 119 × 30 = 3, 570 for *I* = 10. *I* = 14 has a count of 47 × 30 = 1, 410. The 153 × 30 = 4, 590 combinations with a large number of distinct alleles (*I* > 14) are not shown. The theoretical values of 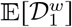 calculated by Eq. 1 from the allele frequencies in the data are visualized in violin plots alongside the theoretical bounds from Figures 1 in Figure 6A. Violin plots for the theoretical 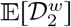 (Eq. 2), empirical 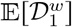, and empirical 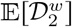, are presented in the remaining panels in a similar manner.

**Figure 6:**
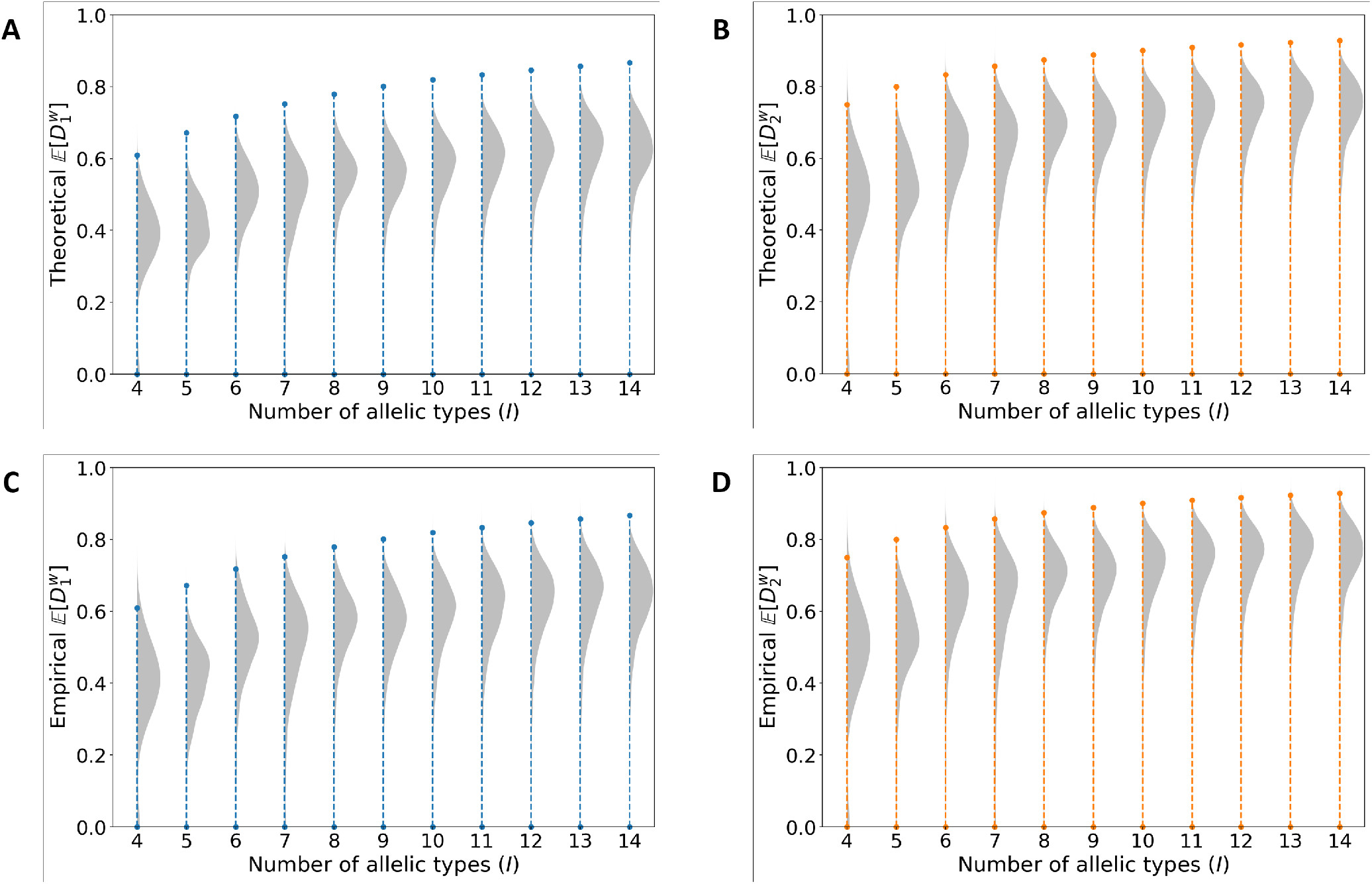
Violin plots of 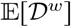 in human population-genetic data; 30 populations with sample size larger than 15 are considered at 630 loci, so that each panel contains 30×630 = 18, 900 data points. Population–locus combinations are grouped by values of *I*; loci with *I* > 14 are not shown. Only half of each violin is depicted. (A) Theoretical 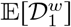 (Eq. 1). (B) Theoretical 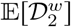 (Eq. 2). (C) Empirical 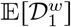. (D) Empirical 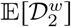. “Theoretical” values are calculated based on the allele frequencies in a population, and “empirical” values are obtained by averaging across all pairs of individuals in the population. The permissible regions for 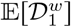 for arbitrary allele frequencies (Theorem 3.2) appear in the background in panels (A) and (C); the permissible regions for 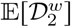 (Theorem 3.6) appear in panels (B) and (D).

The theoretical 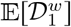 values of populations in the dataset strictly adhere to the mathematical bounds we derived for each *I* (Figure 6A). Similarly, the theoretical 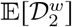 values also adhere to the mathematical bounds (Figure 6B). Data points are concentrated toward the upper bound, a value that can lie substantially below 1. For the empirical 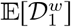 and 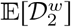, computed from empirical pairwise comparisons of diploid individuals rather than from allele frequencies, the plots are similar (Figure 6C and D). For the empirical values, it is not required that a population–locus computation produce a dissimilarity that lies below the upper bound; nevertheless, nearly all data points do lie below the upper bound (18, 896/18, 900 for 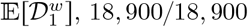 for 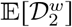).

With the largest allele frequency *M* held fixed, we illustrate the theoretical and empirical dissimilarities in relation to *M* for the case of *I* = 6 (300 population-locus combinations) in Figure 7. The theoretical dissimilarities strictly reside within the permissible region, tending to fill the space toward the upper bound (Figure 7A and B). The empirical dissimilarities generally lie within the permissible region, sometimes extending beyond it (Figure 7C and D). Data points for other values of *I* follow similar patterns.

**Figure 7:**
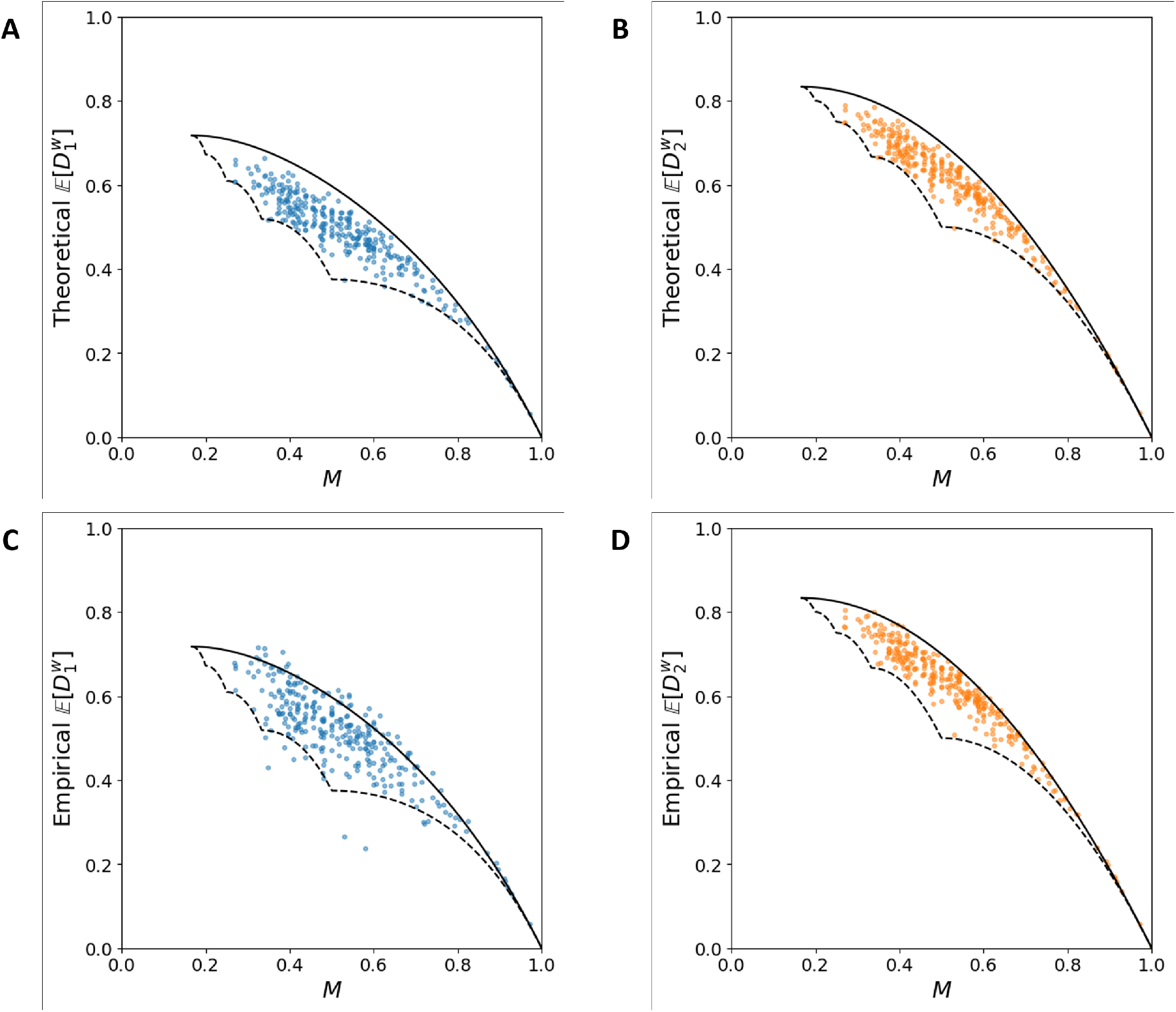
The relationship between the within-population allele-sharing dissimilarity 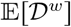 and the largest allele frequency *M* in empirical data. The plot considers 30 populations with sample size larger than 15 and 10 loci with a number of distinct alleles equal to 6, a total of 30 × 10 = 300 population–locus combinations. (A) Theoretical 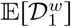 (Eq. 1). (B) Theoretical 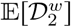 (Eq. 2). (C) Empirical 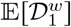. (D) Empirical 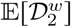. “Theoretical” values are calculated based on the allele frequencies in a population, and “empirical” values are obtained by averaging across all pairs of individuals in the population. The permissible region for 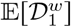 in relation to *M* (Theorem 3.3) appears in panels (A) and (C); the permissible region for 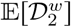 in relation to *M* (Theorem 3.7) appears in panels (B) and (D).

### 5.3 Between-population dissimilarities

We next calculate the theoretical and empirical 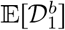 and 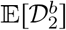 for pairs of populations. Among the 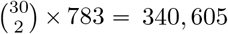 combinations of population pairs and loci, two populations share the most frequent allelic type in 169, 970 (49.9%). We consider these pairs, visualizing the bounds from Theorem 4.2 and Corollary 4.6, which provide the bounds in the scenario in which the two populations in a pair share the same most frequent allelic type at a locus.

The theoretical 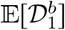 is calculated by determining the allele frequencies of each population and then applying Eq. 3. For the empirical 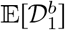, we tabulate pairs of individuals, one from each population. We then compute their 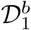 dissimilarities and take an average across all pairs. The process for 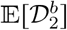 is similar, using Eq. 4. The outcomes for all 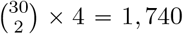 combinations for the 4 loci with *I* = 5 are considered in Figure 8 in a three-dimensional space, showing the 1,092 for which the most frequent allelic type is the same in the two populations and ordering population pairs so that *M*_1_ ⩾ *M*_2_. Comparable patterns are observed for other values of *I*.

**Figure 8:**
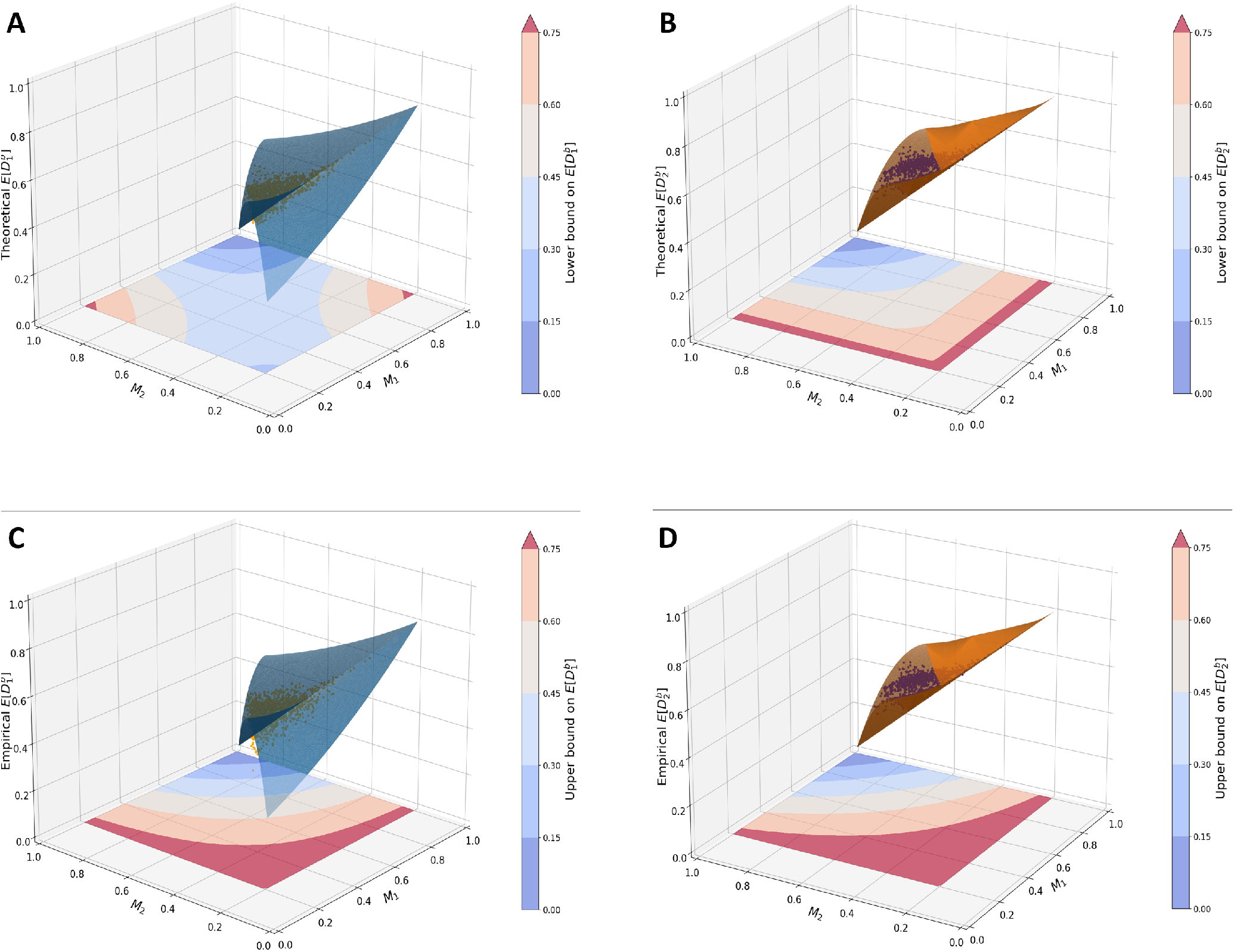
The relationship between the between-population allele-sharing dissimilarity 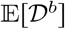 and the largest allele frequencies *M*_1_ and *M*_2_ of two populations that have the same most frequent allelic type. The plot considers 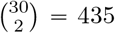 pairs of populations, both with sample size larger than 15, and 4 loci with number of distinct alleles equal to 5, showing the 1,092 of 1,740 combinations for which the two populations have the same most frequent allelic type. Contour plots of the lower and upper bounds are shown in the xy-plane; contour plots cover the whole plane, but for visual simplicity, only the triangle with *M*_1_ ⩾ *M*_2_ is plotted on the z axis. (A) Theoretical 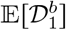 (Eq. 3) and contour plot of the lower bound. (B) Theoretical 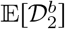 (Eq. 4) and contour plot of the lower bound. (C) Empirical 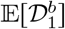 and contour plot of the upper bound. (D) Empirical 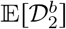 and contour plot of the upper bound. “Theoretical” values are calculated based on the allele frequencies in two populations, and “empirical” values are obtained by averaging across all pairs of individuals, one each from two populations. Each pair of populations is ordered such that *M*_1_ ⩾ *M*_2_. The permissible region for 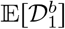 in relation to *M*_1_ and *M*_2_ (Theorem 4.2) appears in panels (A) and (C); the permissible region for 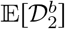 in relation to *M*_1_ and *M*_2_ (Corollary 4.6) appears in panels (B) and (D).

As seen in the data analysis for within-population dissimilarities, the theoretical 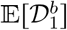 and 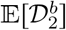 lie strictly within the space bounded by the upper and lower bounds (Figure 8A and B). Note that because we only have loose bounds for 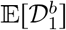, more space exists between the data points representing the theoretical 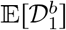 values and the mathematical bounds. 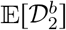 is bounded more tightly. For the empirical 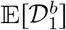 and 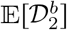, some points fall outside the space demarcated by the bounds (Figure 8C and D).

For the more general case, in which two populations need not have the same allelic type for the most frequent allele, we illustrate the bounds obtained in Theorems 4.3 and 4.5 for all 340,605 combinations of a population pair and locus. The theoretical and empirical 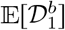 are computed as before. Results for all 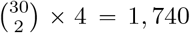 combinations with *I* = 5 appear in Figure 9.

**Figure 9:**
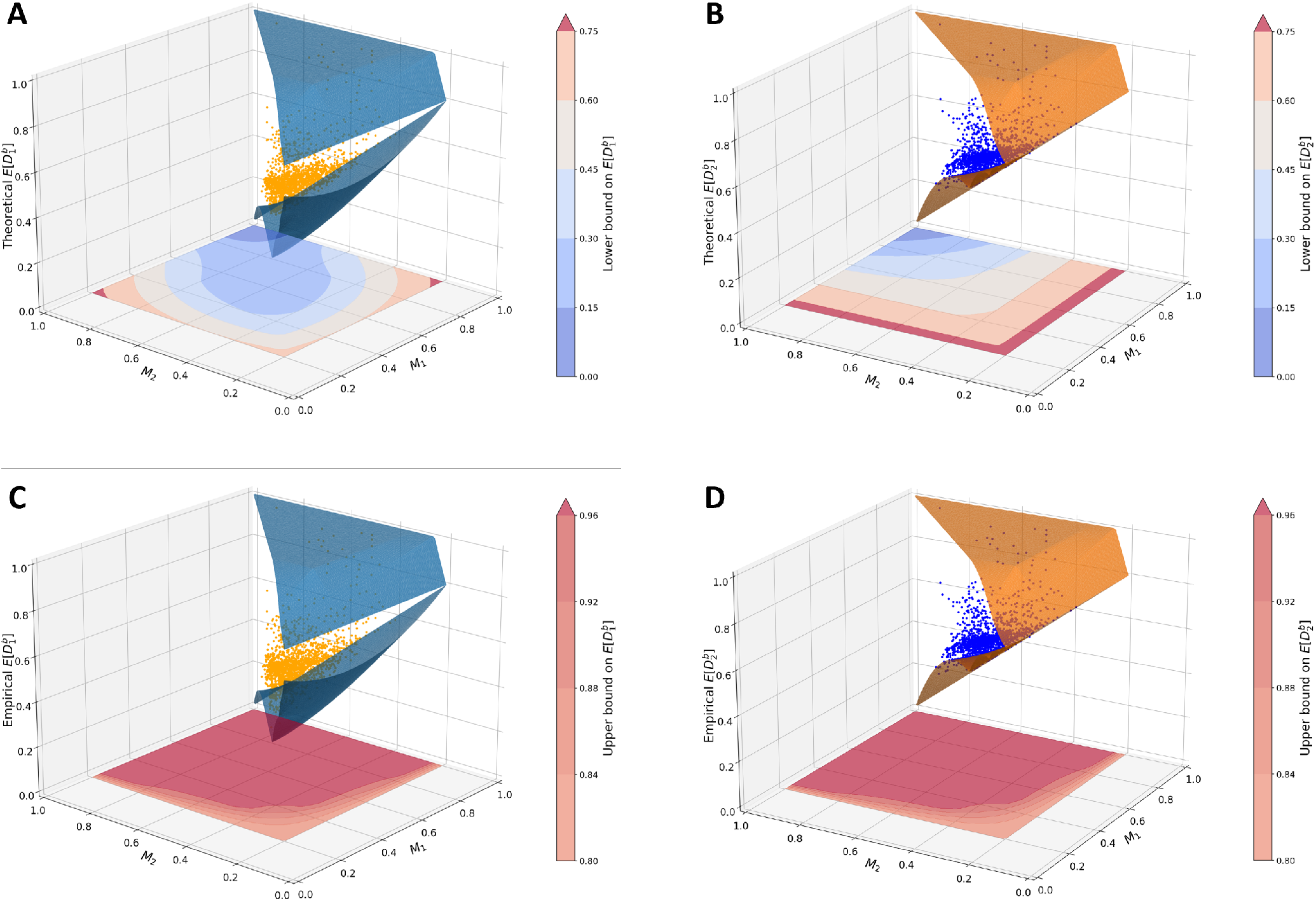
The relationship between the between-population allele-sharing dissimilarity 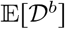 and the largest allele frequencies *M*_1_ and *M*_2_ of two populations that do not necessarily have the same most frequent allelic type. The plot considers 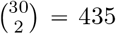 pairs of populations, both with sample size larger than 15, and 4 loci with number of distinct alelles equal to 5, a total of 1740 combinations. Contour plots of the lower and upper bounds are shown in the xy-plane; contour plots cover the whole plane, but for visual simplicity, only the triangle with *M*_1_ ⩾ *M*_2_ is plotted on the z axis. (A) Theoretical 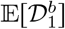 (Eq. 3) and contour plot of the lower bound. (B) Theoretical 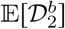 (Eq. 4) and contour plot of the lower bound. (C) Empirical 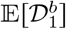 and contour plot of the upper bound. (D) Empirical 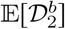 and contour plot of the upper bound. “Theoretical” values are calculated based on the allele frequencies in two populations, and “empirical” values are obtained by averaging across all pairs of individuals, one each from two populations. Each pair of populations is ordered such that *M*_1_ ⩾ *M*_2_. The permissible region for 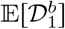 in relation to *M*_1_ and *M*_2_ (Theorem 4.3) appears in panels (A) and (C); the permissible region for 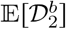 in relation to *M*_1_ and *M*_2_ (Theorem 4.5) appears in panels (B) and (D).

Most of the theoretical dissimilarities congregate within the central area of the permissible region (Figure 9A and B). The permissible region is generally larger than in the case in which the most frequent allelic type is the same for a pair of populations, as seen in Figure 8. In the case of *I* = 5, the empirical mean dissimilarities all fall within the permissible range (Figure 9C and D).

### 5.4 Allele-sharing dissimilarity and heterozygosity

A notable property of 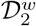 is that the expression for its expectation is exactly identical to the expression 1−*σ*_2_ for the heterozygosity of a population, as computed from its allele frequencies. We compare the theoretical and empirical allele-sharing dissimilarities to heterozygosity in two ways. First, we compute the theoretical heterozygosity for each of the 7 geographic regions; this quantity is precisely 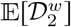 for those regions. Next, we compute the theoretical heterozygosity for each of the 53 sampled populations, the value of 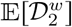 for the populations.

Figure 10A plots the theoretical 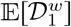 and 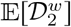 in relation to the theoretical heterozygosity 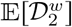 at the regional level, showing 7 × 783 = 5, 481 points. The values of 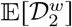 follow the *y* = *x* line, as *x* and *y* values are equal. The values of 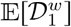 lie below the *y* = *x* line, in accord with Theorem 4.3, which—by specifying that two populations have identical frequencies—can be seen to demonstrate that the theoretical 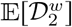 provides an upper bound for the theoretical 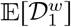. Similar results are obtained in Figure 10B for the 53 × 783 = 41, 499 data points at the population level.

**Figure 10:**
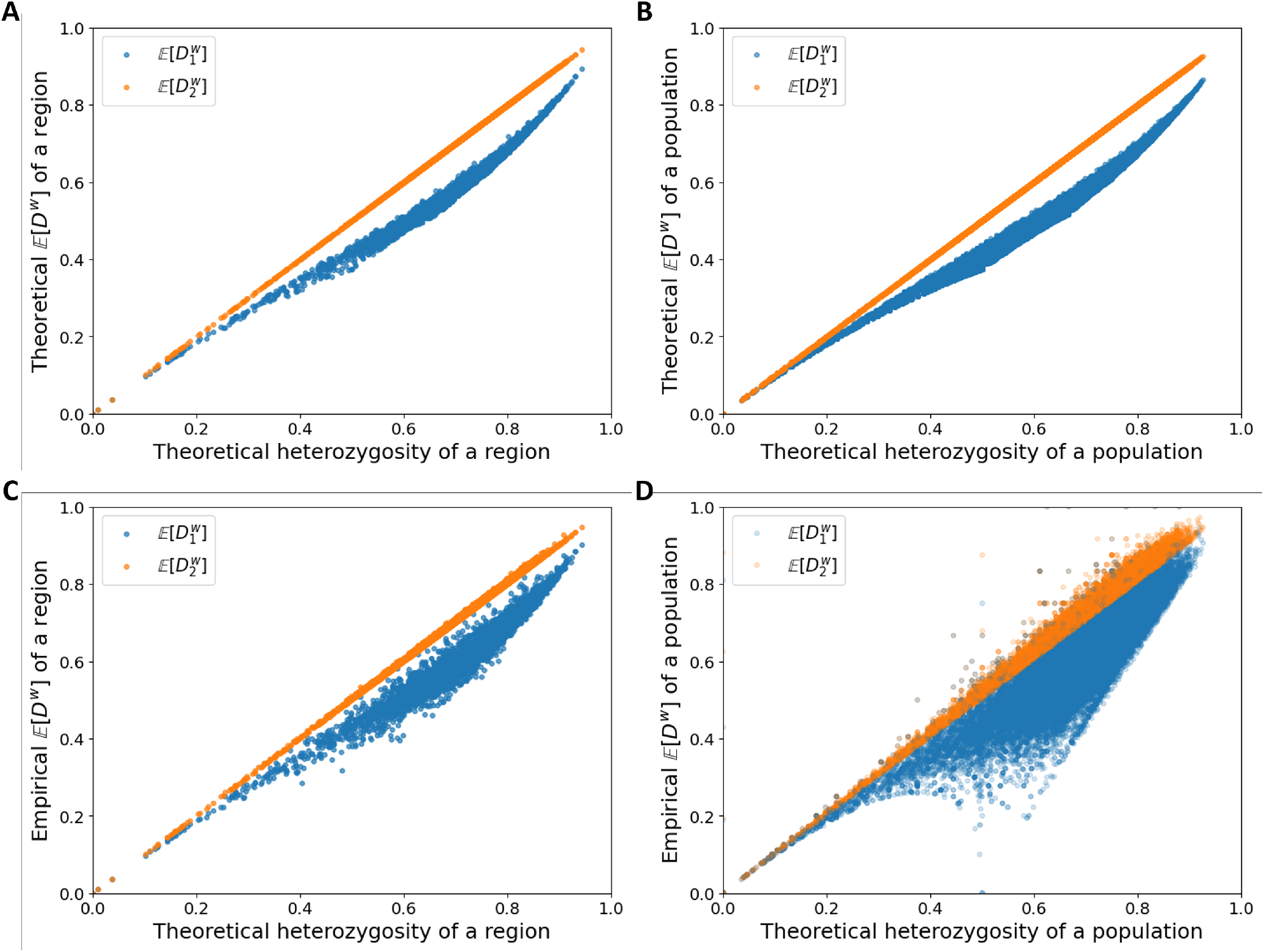
Within-population allele-sharing dissimilarity in relation to theoretical heterozygosity 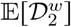. The plots consider 7 geographic regions and 53 populations, for a total of 7 × 783 = 5, 481 points in panels (A) and (B), and 53 × 783 = 41, 499 points in panels (C) and (D) (A) Theoretical 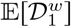 and 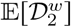 for regions (Eqs. 1 and 2). (B) Theoretical 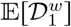 and 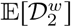 for populations (Eqs. 1 and 2). (C) Empirical 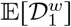 and 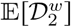 for regions. (D) Empirical 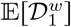 and 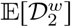 for populations. “Theoretical” values are calculated based on the allele frequencies in a population, and “empirical” values are obtained by averaging across all pairs of individuals in the population.

Next, we examine the empirical 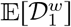 and 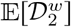 in relation to the theoretical heterozygosity 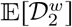, computing allele-sharing dissimilarity by considering pairs of individuals in a region or population. Figure 10C plots the 7 × 783 = 5, 481 data points at the regional level, and Figure 10D plots the 53 × 783 = 41, 499 data points at the population level. In both panels, the empirical 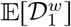 values are more variable than the 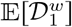 values in Figure 10A and B. The empirical 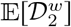 values do not precisely equal to the theoretical 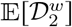 values, though the empirical and theoretical values are quite similar.

### 5.5 Mathematical bounds in empirical allele-sharing dissimilarities

Visualizations of distributions of pairwise genetic dissimilarities between individuals have been important for understanding empirical genetic differences, notably in human populations [Mountain and Ramakrishnan, 2005, Rosenberg, 2011]. In Figure 5 of Rosenberg [2011], distributions of pairwise genetic dissimilarities between individuals, as computed by 𝒟_1_, are presented in various computations within regions and within geographic regions.

We reproduce Figure 5B and C of Rosenberg [2011], illustrating how the distributions of empirical genetic dissimilarities are informed by mathematical bounds. The calculation uses all 1,048 individuals and 53 populations in the data. In Figure 11A, we show the empirical distribution of allele-sharing dissimilarity between pairs of individuals within regions, averaging across all 783 loci and replotting Figure 5B of Rosenberg [2011]. In Figure 11B-H, we show the empirical distributions within single regions, plotting them alongside mathematical bounds on 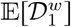 for the region. The bounds are calculated from the region-wise allele frequencies for a locus according to Theorem 3.3, then averaged across all loci to obtain the mean lower and upper bounds.

**Figure 11:**
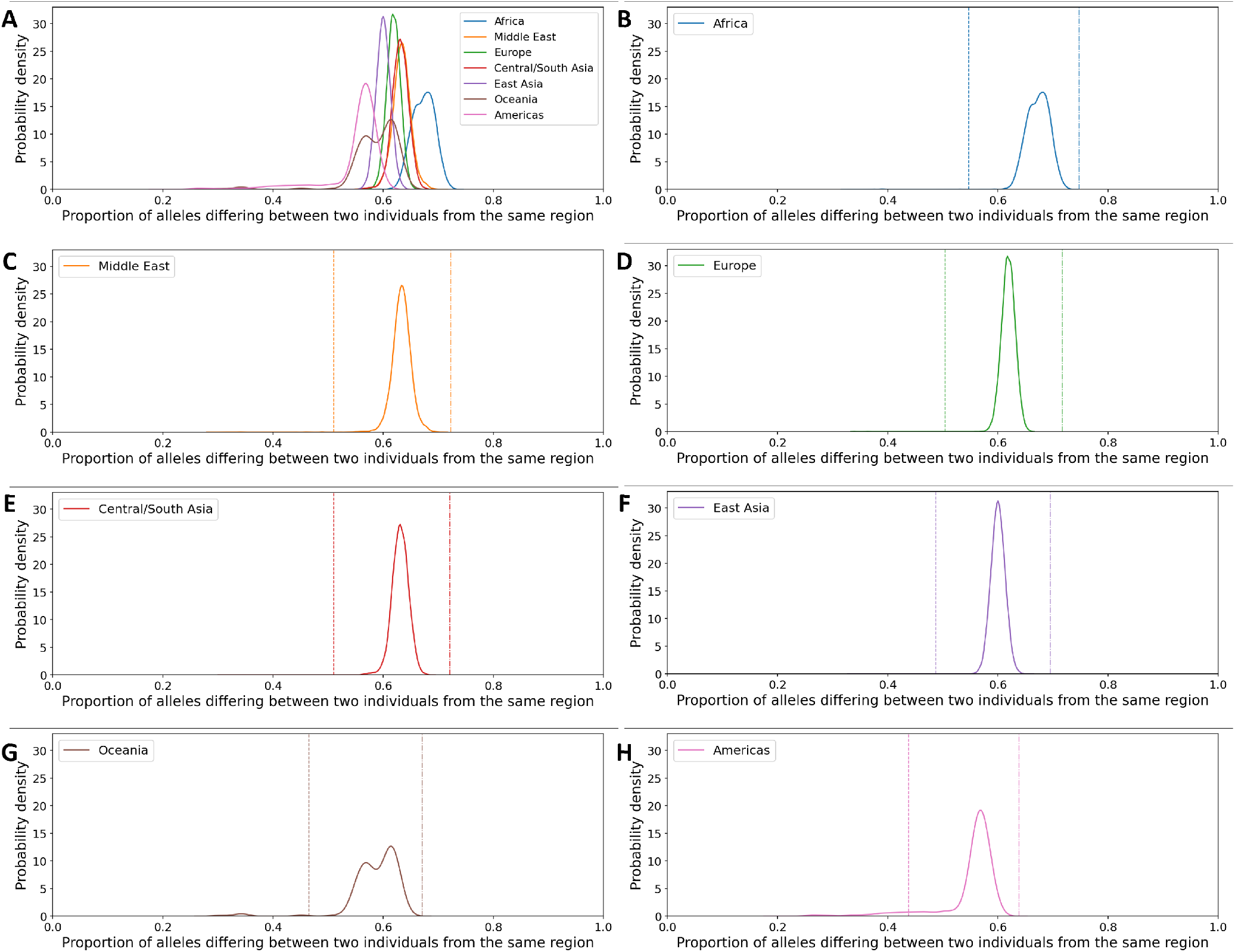
Distributions of empirical values of within-region 𝒟_1_ averaged across all 783 loci for pairs of individuals in human population-genetic data. For each region, mathematical bounds for 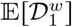 are calculated from allele frequencies within a region according to Theorem 3.3, averaging across loci. (A) Seven regions displayed together. (B) Africa. (C) Middle East. (D) Europe. (E) Central/South Asia. (F) East Asia. (G) Oceania. (H) Americas.

In Figure 12A, we similarly show the empirical distribution of allele-sharing dissimilarity between pairs of individuals within populations, averaging across all 783 loci and replotting Figure 5C of Rosenberg [2011]. In Figure 12B-H, we show the empirical distributions of within-population dissimilarities grouped by region, plotting them alongside mathematical bounds on 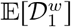 for single regions. The bounds are calculated from population-wise allele frequencies for a locus via Theorem 3.3, then averaged across populations within a region and then across all loci to obtain the mean bounds for a region.

**Figure 12:**
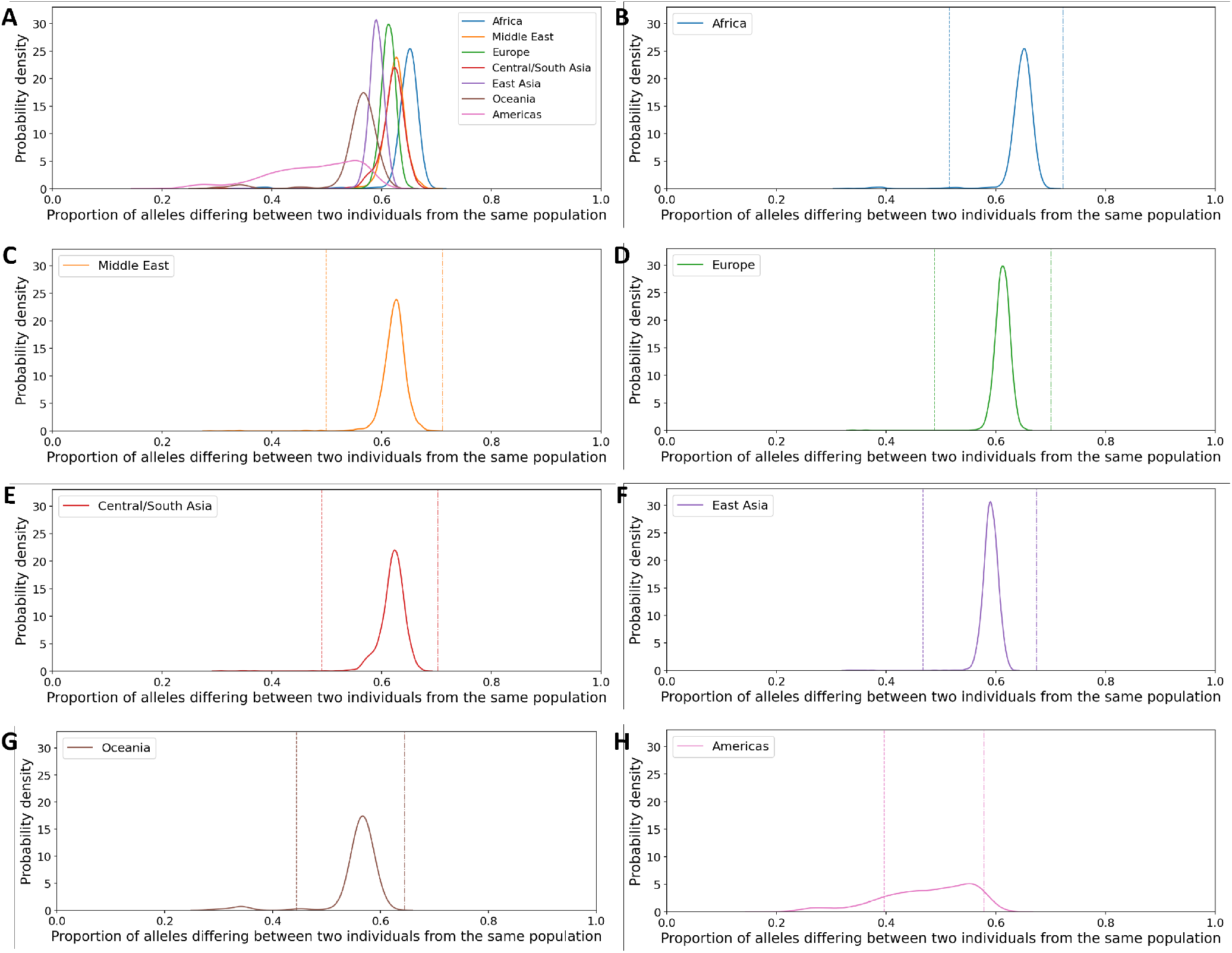
Distributions of empirical values of within-population 𝒟_1_ averaged across all 783 loci for pairs of individuals in human population-genetic data. For each region, mathematical bounds for 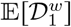 are calculated for each population from allele frequencies within the population according to Theorem 3.3, averaging across loci. Bounds are then averaged across populations within regions. (A) Seven regions displayed together. (B) Africa. (C) Middle East. (D) Europe. (E) Central/South Asia. (F) East Asia. (G) Oceania. (H) Americas.

Both in Figure 11 and in Figure 12, the theorem specifies a relatively narrow range for values of 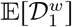, dependent on the particular values of the frequency *M* of the most frequent allelic type in the empirical data. Most of the probability mass lies between the lower and upper bounds. Some empirical dissimilarity values lie outside the range specified by the bounds; it is not required that an empirical dissimilarity lie between the bounds, as the bounds are obtained from an average of theoretical values across loci, whereas the empirical values are obtained for pairs of individuals. Nevertheless, the plots suggest that the mathematical bounds specify informal constraints on the distribution of empirical values of the allele-sharing dissimilarity in population-genetic data.

## 6 Discussion

Allele-sharing dissimilarities, computed theoretically as expectations based on allele-frequency distributions or empirically based on pairs of individuals, have often been used for studying genetic variation in populations. We have shown that as a function of properties of allele-frequency distributions, the range for expected allele-sharing dissimilarities is substantially narrower than the unit interval. Specifically, considering dissimilarities 𝒟_1_ and 𝒟_2_, we have obtained mathematical expressions for constraints on expected ASD within a population when the number of allelic types is fixed (Theorems 3.2 and 3.6), as well as when the frequency of the most frequent allelic type is also fixed (Theorems 3.3 and 3.7). Additional mathematical results concern the area of the region bounded between the smallest and largest within-population ASD values as a function of number of distinct alleles. This region increases in size with an increasing number of allelic types, converging to a value well below 1 (Propositions 3.4 and 3.8). We have also obtained corresponding expressions in between-population scenarios with the number of allelic types fixed (Propositions 4.1 and 4.4) and additionally with fixed frequencies for the most frequent allelic type (Theorems 4.2, 4.3, and 4.5, and Corollary 4.6).

In illustrations of the mathematical results using data from human populations, we have found that empirical mean ASD values reflect the theoretical expectations computed from allele-frequency distributions (Figures 6-10). The mathematical bounds on ASD values in relation to the frequency of the most frequent allelic type suggest that ASD values are expected to vary in relatively narrow ranges within the unit interval; indeed, empirical distributions of 𝒟_1_ are quite constrained (Figures 11 and 12). The mathematical results assist in explaining the relatively narrow ranges for ASD values computed in worldwide human populations, as the frequency of the most frequent allelic type constrains the between-population ASD values.

The bounds are meaningful beyond these computations. In particular, in between-population analyses, a larger range between the bounds permits more variability in the dissimilarity across pairs of populations. Such variability can be relevant in applications that rely on distinguishing the ASD values for different pairs of groups, as greater variability indicates a greater potential to distinguish values for different pairs. Theoretical properties of methods such as neighbor-joining tree construction and multidimensional scaling that rely on dissimilarity matrices, and the effects on these methods of the range between the bounds, can be explored more specifically.

We have considered two ASD measures, 𝒟_1_, which was used in the data example mimicking the analysis of Rosenberg [2011] (Figures 11 and 12), and 𝒟_2_, which provides a generalization of heterozygosity (Figure 10). For within-population computations, bounds are provided for both dissimilarities. For between-population computations, however, for 𝒟_1_, mathematical analysis is more limited. Owing to simpler mathematical expressions, tight bounds can be obtained for 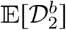 in the between-population case. For 𝒟_1_, mathematical bounds in Theorem 4.2 are loose in the case that 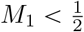 or 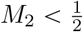.

Limitations of the study include the fact that the constraints on the expected allele-sharing dissimilarity consider only on the most frequent allelic type. The frequencies of subsequent allelic types might impose constraints that might be of interest for future investigation, as occurs in various other contexts [Garud and Rosenberg, 2015, Morrison and Rosenberg, 2023]. We also note that in our empirical analysis, we average across all pairs of individuals, either within or between populations, to obtain the empirical 𝔼[𝒟]. The reuse of each individual in multiple pairs violates the assumption that pairs are independent draws from the allele-frequency distributions, so that the empirical results do not quite mimic the computation performed theoretically. The theoretical results assume that pairs of alleles within an individual are independently drawn from the allele-frequency distribution— but empirically, the two alleles can be dependent due to inbreeding. The violation of the assumptions can contribute to deviations of the empirical observations from the theoretical values.

Additionally, our mathematical expressions are for dissimilarity values computed based on a single genetic locus. In empirical studies such as Rosenberg [2011], however, measures are typically calculated on multiple loci and averaged together. An explicitly multilocus analysis that considers the constraints at multiple loci could provide further insight into the behavior of an empirical mean across many loci.

## Acknowledgments

We acknowledge National Institutes of Health grant R01 HG005855 for support.

## A. Proof of Lemma 3.1

*f* (**p**) = 1 − 2*σ*_2_ + 2*σ*_3_ − *σ*_4_ is symmetric in the *p*_*i*_ by construction, and all first partial derivatives 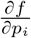 exist. By Theorem 2.3, to show that *f* is Schur-concave, it suffices to show that 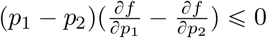 for all **p**∈ Δ^*I*−1^.

We have 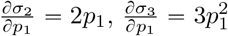, and 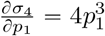, so that

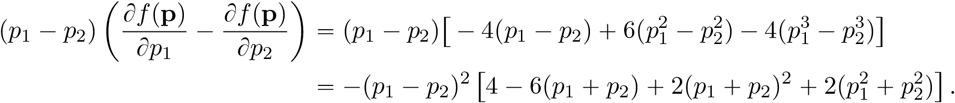

For 0 ⩽ *x* ⩽ 1, 4 − 6*x* + 2*x*^2^ ⩾ 0 with equality if and only if *x* = 1. Hence 4 − 6(*p*_1_ + *p*_2_) + 2(*p*_1_ + *p*_2_)^2^ ⩾ 0 always holds for 0 ⩽ *p*_1_ + *p*_2_ ⩽ 1. We then have 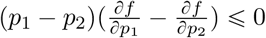. By Theorem 2.3, *f* is Schur-concave.

To verify strict Schur-concavity, note that 4 − 6(*p*_1_ + *p*_2_) + 2(*p*_1_ + *p*_2_)^2^ = 0 requires *p*_1_ + *p*_2_ = 1, so that 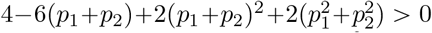 for all permissible (*p*_1_, *p*_2_): either *p*_1_+*p*_2_ ≠ 1 and 4−6(*p*_1_+*p*_2_)+2(*p*_1_+*p*_2_)^2^ > 0, or *p*_1_ + *p*_2_ = 1 and 2(*p*_1_ + *p*_2_)^2^ > 0. We conclude that 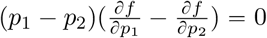 implies *p*_1_ = *p*_2_.

## B Proof of Proposition 3.4

The desired area is calculated by considering *M* in segments. For 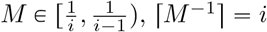. The area then equals

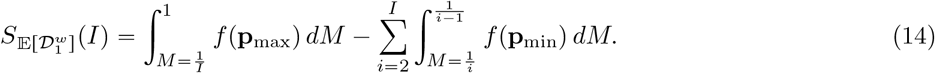

The first term is

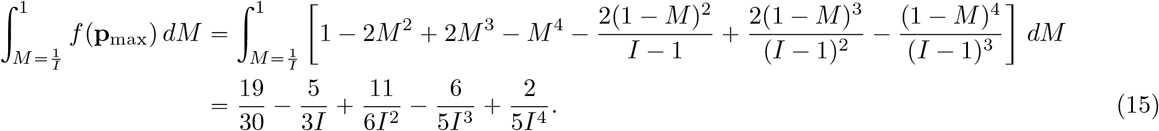

The second term is

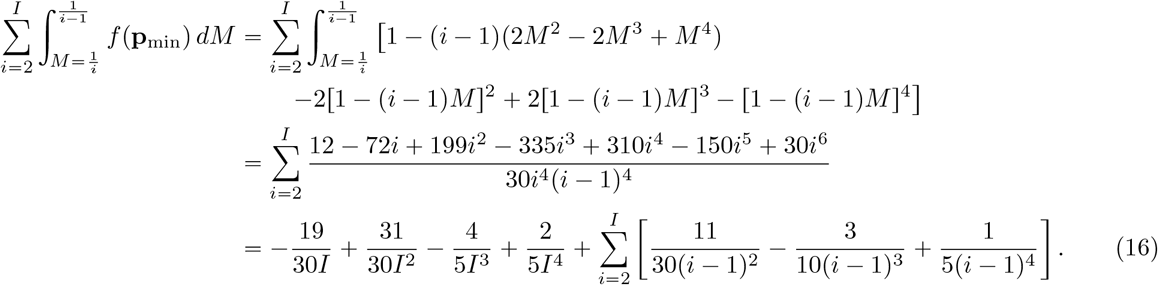

Subtracting Eq. 16 from Eq. 15 in Eq. 14, we obtain the quantity in Eq. 5.

## C Proof of Theorem 4.2

First, for the upper bound, because 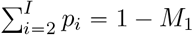 and 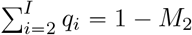, Eq. 3 can be written

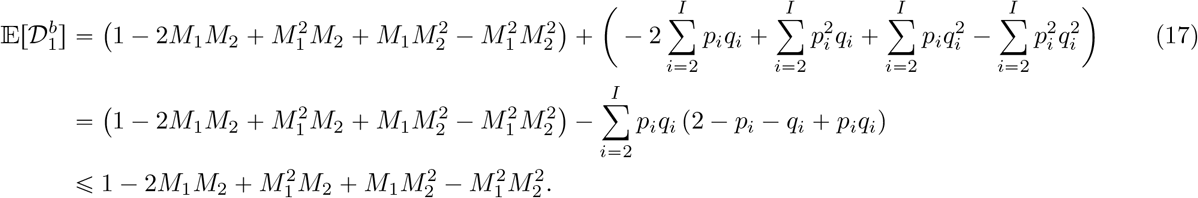

The last inequality holds because 0 ⩽ *p*_*i*_ < 1 and 0 ⩽ *q*_*i*_ < 1 for all *i* = 2, 3, …, *I*, so that 2 − *p*_*i*_ − *q*_*i*_ + *p*_*i*_*q*_*i*_ > 0, *p*_*i*_*q*_*i*_ ⩾ 0, and 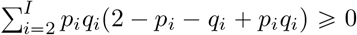. Equality with the upper bound requires that for all *i*, 2 ⩽ *i* ⩽ *I, p*_*i*_*q*_*i*_ = 0. That is, for all *i* = 2, 3, …, *I, p*_*i*_ = 0 or *q*_*i*_ = 0, so that allele 1 is the only allele shared between populations.

Next, for the lower bound,

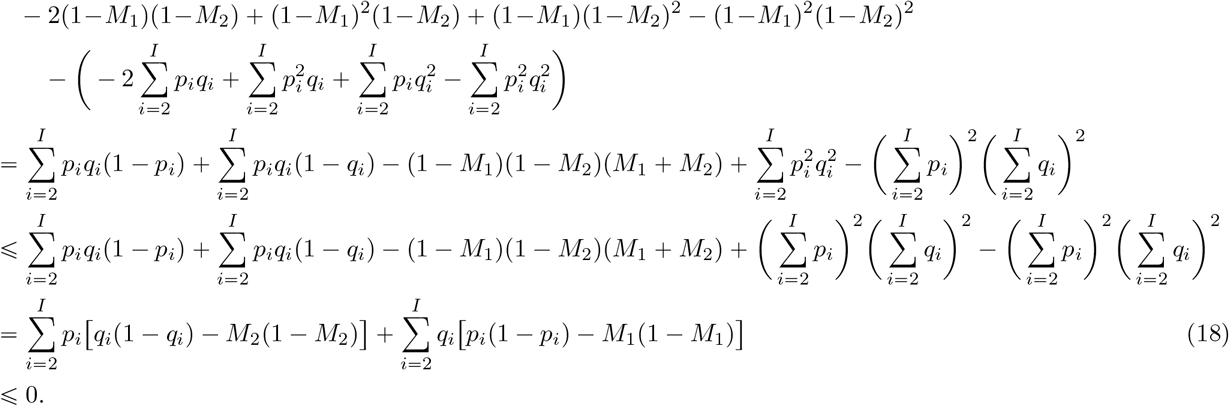

The first inequality uses the fact that the *p*_*i*_ and *q*_*i*_ are all non-negative, so that 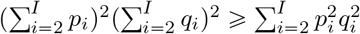. The last inequality uses the fact that *p*_*i*_ ⩽ *M*_1_, *p*_*i*_ ⩽ 1−*M*_1_, *q*_*i*_ ⩽ *M*_2_, and *q*_*i*_ ⩽ 1−*M*_2_. The function *f* (*x*) = *x*(1−*x*) is nondecreasing for 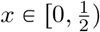, and one of *M*_1_ and 1 − *M*_1_ must lie in 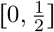, so *p*_*i*_ ⩽ *M*_1_ and *p*_*i*_ ⩽ 1 − *M*_1_ implies *f* (*p*_*i*_) ⩽ *f* (*M*_1_); analogously, *f* (*q*_*i*_) ⩽ *f* (*M*_2_).

Applying Eq. 17, we therefore have

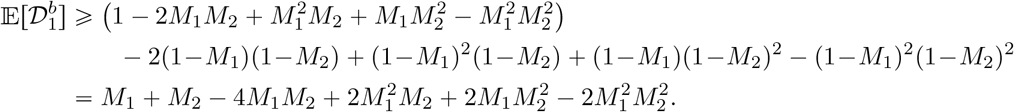

Equality with the lower bound requires 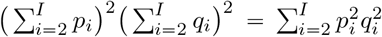. Because *p*_*2*_⩾ *p*_*3*_⩾ … ⩾ *p*_*I*_, this condition requires *p*_2_ = 1 − *p*_1_ = 1 − *M*_1_ and hence *q*_2_ = 1 − *q*_1_ = 1 − *M*_2_, making use of assumptions 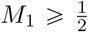 and 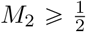_2 2_. Equality with the lower bound also requires that the expression in Eq. 18 equal 0; allele-frequency distributions (*p*_1_, *p*_2_, *p*_3_, …, *p*_*I*_) = (*M*_1_, 1 − *M*_1_, 0, …, 0) and (*q*_1_, *q*_2_, *q*_3_, …, *q*_*I*_) = (*M*_2_, 1 − *M*_2_, 0, …, 0) produce a value of 0 in Eq. 18.

## D Proof of Theorem 4.3

We write 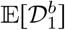 in the form

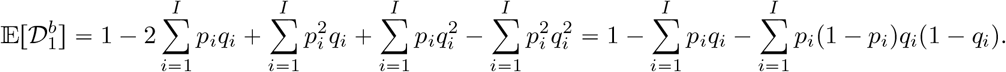

For the upper bound, because 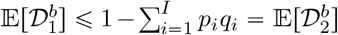, the upper bound of 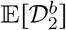 can also serve as a (loose) upper bound for 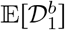.

To obtain a loose lower bound, we must bound from above the quantity 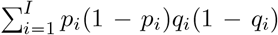 given max{*p*_1_, *p*_2_, …, *p*_*I*_} = *M*_1_ and max{*q*_1_, *q*_2_, …, *q*_*I*_} = *M*_2_. First, note that for *p*_1_, *p*_2_, …, *p*_*I*_ with *p*_1_ ⩾ *p*_2_ ⩾ … *p*_*I*_ ⩾ 0 and 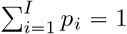, we have *p*_*i*_(1 −*p*_*i*_) ⩾ *p*_*j*_(1 −*p*_*j*_) for *i* < *j*. This result follows because *f* (*x*) = *x*(1 −*x*) is maximized at 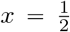, declining symmetrically around the maximum, and *p*_*j*_ lies farther from 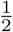 than does *p*_*i*_; the claim is verified in two cases, 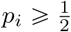, for which 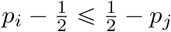, and 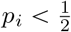, for which 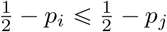.

It follows that for each *i, p*_*i*_(1−*p*_*i*_) ⩽ *M*_1_(1−*M*_1_) and *q*_*i*_(1−*q*_*i*_) ⩽ *M*_2_(1−*M*_2_), so that 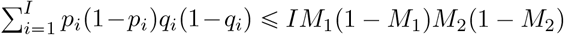.

## E Proof of Theorem 4.5

Theorem 4.5 states the bounds on 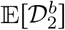 and gives sufficient conditions on the *p*_*i*_ and *q*_*i*_ at which the bounds are reached—given an upper bound *M*_1_ on the *p*_*i*_ and an upper bound *M*_2_ on the *q*_*i*_ (also assuming 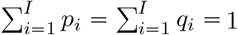). The upper bounds need not occur at the same allele.

The proof proceeds by a series of lemmas. Informally, Lemma E.1 shows that for fixed *a*_*i*_, we can reduce the sum of products 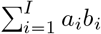 by a particular choice of the value of a specific frequency *b*_*ℓ*_ (if it is not already optimized).

### Lemma E.1.

*Suppose a collection of I* ⩾ 2 *fixed non-negative values a*_1_, *a*_2_, …, *a*_*I*_ *is given, with a*_1_ ⩾ *a*_2_ ⩾ … ⩾ *a*_*I*_. *Suppose b*_1_, *b*_2_, …, *b*_*I*_ *are non-negative values satisfying three conditions:*

1. *monotonicity, b*_1_ ⩽ *b*_2_ ⩽ … ⩽ *b*_*I*_;
2. *fixed total sum*, 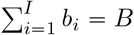; *and*
3. *boundedness from above, b*_*i*_ ⩽ *b*\* *for all i* = 1, 2, …, *I, where* 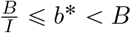.

*Consider ℓ with* 2 ⩽ *ℓ* ⩽ *I and* (*I* − *ℓ*)*b*\* < *B. Suppose b*_*i*_ = *b*\* *for each i with ℓ* < *i* ⩽ *I, and suppose b*_*ℓ*_ < min (*b*\*, *B* − (*I* − *ℓ*)*b*\*). *Then there exists a set of values* 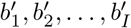 *with* 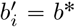 *for each i with l* < *i ⩽ I, satisfying conditions* (*1*), (*2*), *and* (*3*), *such that b*^1^ = min *b*\*, *B* −(*I* − *ℓ*)*b*\*) and

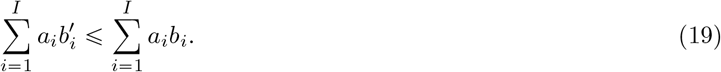

*Proof*. For convenience, write *s* = min (*b*\*, *B* −(*I* −*ℓ*)*b*\*), so that 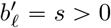. Let 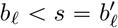. We have 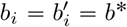 for each *i* with *ℓ* < *i* ⩽ *I*. 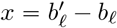, a positive quantity representing the difference between the value we will place in the *ℓ*th entry in our new sequence and the value in the current sequence. Because 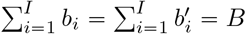,

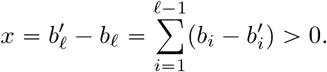

Let *k* be the unique index that satisfies 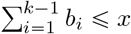 and 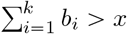. We set the values of 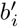 so that 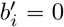 for each *i* with 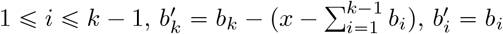 for *k* + 1 ⩽ *i* ⩽ *ℓ* − 1, and 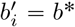 for *ℓ* ⩽ *i* ⩽ *I*.

Note that *k* ⩽ *ℓ* always holds. For contradiction, suppose *k* > *ℓ*. Then 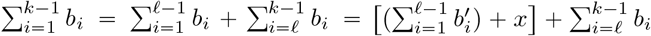. We have 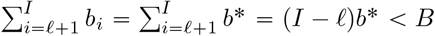; because 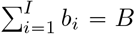, it follows that 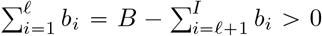. Next, because *b*_*ℓ*_ ⩾ *b*_*i*_ for each *i* with 1 ⩽ *i* ⩽ *ℓ* − 1, we have *b*_*ℓ*_ > 0. As a result, 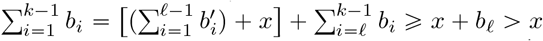 contradicting the condition 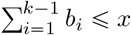 in the definition of *k*.

We have constructed a sequence of values 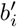 that continues to satisfy the monotonicity, fixed-total-sum, and boundedness-from-above conditions. (1) For monotonicity, 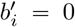 for 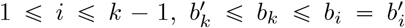 for 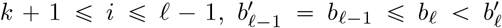, and 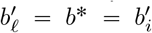 for 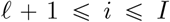. (2) For fixed total sum, 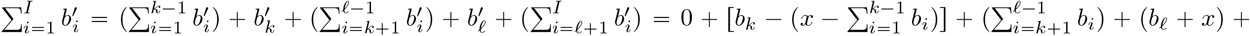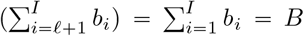. (3) For boundedness from above, 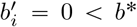 for 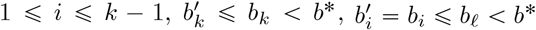 for *k* + 1 ⩽ *i* ⩽ *ℓ* − 1, and 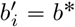 for *ℓ* ⩽ *i* ⩽ *I*.

It remains to show that Eq. 19 holds. We have

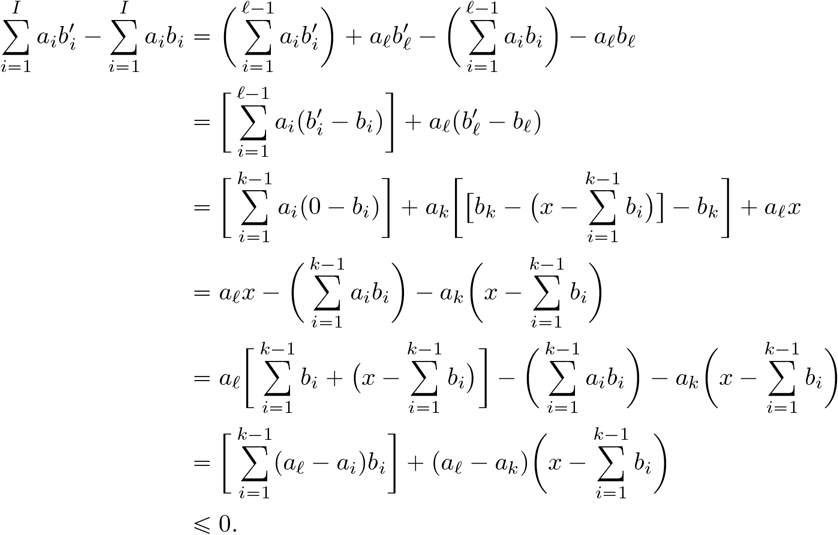

In the last step, the inequality holds because *k* ⩽ *ℓ* and the *a*_*i*_ are monotonically decreasing, so that *a*_*ℓ*_ ⩽ *a*_*i*_ for all *i*, 1 ⩽ *i* ⩽ *ℓ*.

Lemma E.2 is similar to Lemma E.1, but in the reverse direction. It shows that for fixed *a*_*i*_, we can *increase* 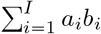 by a particular choice of the value of a specific frequency *b*_*ℓ*_ (if it is not already optimized).

### Lemma E.2.

*Suppose a collection of I* ⩾ 2 *fixed non-negative values a*_1_, *a*_2_, …, *a*_*I*_ *is given, with a*_1_ ⩾ *a*_2_ ⩾ … ⩾ *a*_*I*_. *Suppose b*_1_, *b*_2_, …, *b*_*I*_ *are non-negative values satisfying three conditions:*

1. *monotonicity, b*_1_ ⩾ *b*_2_ ⩾ … ⩾ *b*_*I*_;
2. *fixed total sum*, 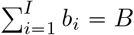; *and*
3. *boundedness from above, b*_*i*_ ⩽ *b*\* *for all i* = 1, 2, …, *I, where* 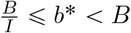.

*Consider ℓ with* 1 ⩽ *ℓ* ⩽ *I* − 1 *and* (*ℓ* − 1)*b*\* < *B. Suppose b*_*i*_ = *b*\* *for each i with* 1 ⩽ *i* < *ℓ, and suppose b*_*ℓ*_ < min(*b*\*, *B* − (*ℓ* − 1)*b*^∗)^. *Then there exists a set of values* 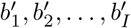 *with* 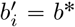 *for each i with* 1 ⩽ *i* < *ℓ, satisfying conditions* (*1*), (*2*), *and* (*3*), *such that* 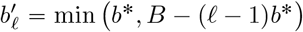, *and*

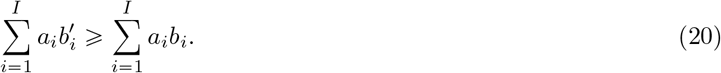

*Proof*. The proof is similar to that of Lemma E.1. Write *s* = min (*b*\*, *B* − (*ℓ* − 1)*b*\*), so that 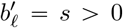. Let 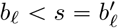. We now have 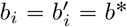 for each *i* with 1 ⩽ *i* < *ℓ*. Let 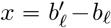 positive quantity representing the difference between the value we place in the *ℓ*th entry in our new sequence and the value in the current sequence. Because 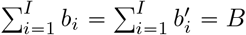,

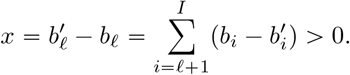

Let *k* be the unique index that satisfies 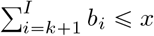 and 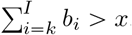. We set the values of 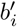 so that 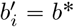 for 1 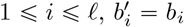 for 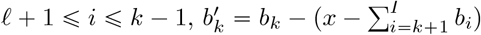, and 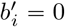 for each *i* with *k* + 1 ⩽ *i* ⩽ *I*.

We show *k* ⩾ *ℓ*. For contradiction, suppose *k* < *ℓ*. Then 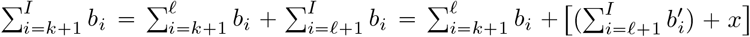. We have 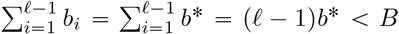; because 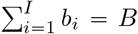, it follows that 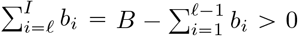. Next, because *b*_*ℓ*_ ⩾ *b*_*i*_ for each *i* with *ℓ* + 1 ⩽ *i* ⩽ *I*, we have *b*_*ℓ*_ > 0. As a result,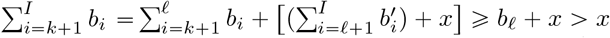, contradicting the condition 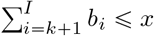 in the definition of *k*.

The constructed sequence of values 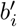 continues to satisfy the monotonicity, fixed-total-sum, and boundedness-from-above conditions. (1) For monotonicity, 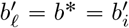 for 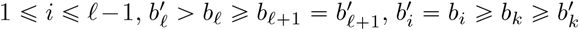 for *ℓ* + 1 ⩽ *i* ⩽ *k* − 1, and 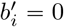 for *k* + 1 ⩽ *i* ⩽ *I*. (2) For fixed total sum, 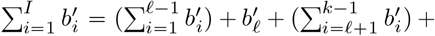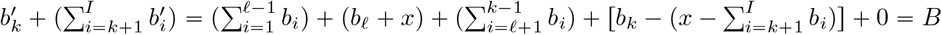. (3) For boundedness from above, 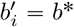 for 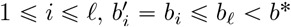 for 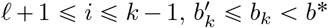, and 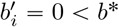 for *k* +1 ⩽ *i* ⩽ *I*.

It remains to show that Eq. 20 holds. We have

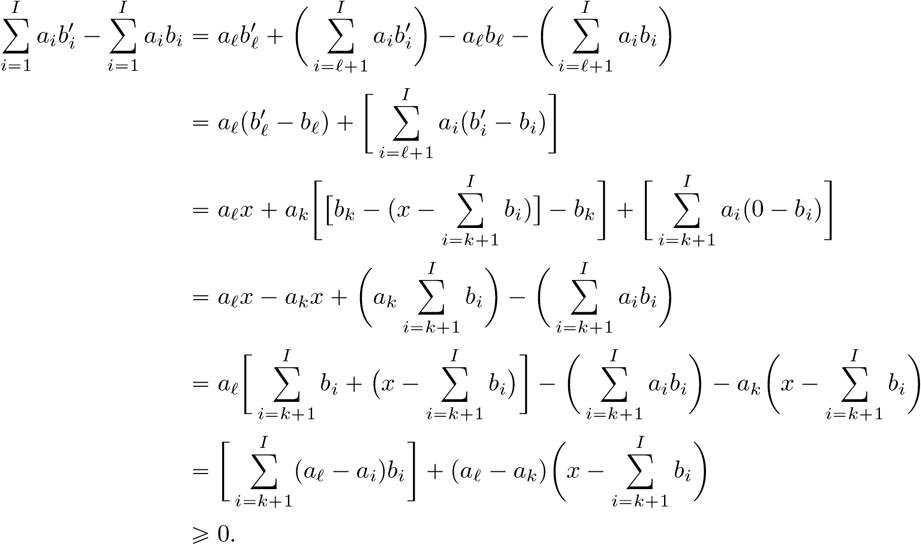

In the last step, the inequality holds because *k* ⩾ *ℓ* and the *a*_*i*_ are monotonically decreasing, so that *a*_*ℓ*_ ⩾ *a*_*i*_ for all, *i*, 𝓁 ⩽ *i* ⩽ *I*.

Lemma E.3 now uses Lemmas E.1 and E.2 to find the minimum and maximum of the sum of products 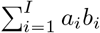, allowing both *a*_*i*_ and *b*_*i*_ to vary.

### Lemma E.3.

*Consider all possible sets of non-negative real numbers* {*a*_1_, *a*_2_, …, *a*_*I*_} *and* {*b*_1_, *b*_2_, …, *b*_*I*_} *with fixed sums* 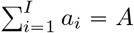 *and* 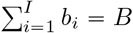, *where I* ⩾ 2, *A* > 0, *and B* > 0. *Suppose that the a*_*i*_ *are non-decreasing, with a*_1_ ⩾ *a*_2_ ⩾ … ⩾ *a*_*I*_ *and that the b*_*i*_ *are monotonic, with b*_1_ ⩾ *b*_2_ ⩾ … ⩾ *b*_*I*_ *or b*_1_ ⩽ *b*_2_ ⩽ … ⩽ *b*_*I*_. *Suppose also that* 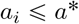 *and* 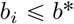 *for all i, with* 0 < *a*\* < *A and* 0 < *b*\* < *B. Let α* = ⌈ *A*/*a*\* ⌉. *and β* = ⌈ *B*/*b*\* ⌉. *The values of I, A, B, a*\*, *and b*\* *are fixed and given. Consider the following conditions:*

1. *a*_*i*_ = *a*\* *for* 1 ⩽ *i* ⩽ *α* − 1, *a*_*α*_ = *A* −(*α* − 1)*a*\*, *and a*_*i*_ = 0 *for α* + 1 ⩽ *i* ⩽ *I*.
2. *b*_*i*_ = 0 *for* 1 ⩽ *i* ⩽ *I* − *β, b*_*I*−*β*1_ = *B* −(*β* − 1)*b*\*, *and b*_*i*_ = *b*\* *for I* − *β* + 2 ⩽ *i* ⩽ *I*.
3. *b*_*i*_ = *b*\* *for* 1 ⩽ *i* ⩽ *β* − 1, *b*_*β*_ = *B* −(*β* − 1)*b*\*, *and b*_*i*_ = 0 *for β* + 1 ⩽ *i* ⩽ *I*.

*Then* (*i*) 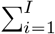 *a*_*i*_*b*_*i*_ *achieves its maximal value if Conditions 1 and 3 hold*. (*ii*) 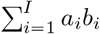 *achieves its minimal value if Conditions 1 and 2 hold*.

*Proof*. (i) For the upper bound, by the rearrangement inequality (Theorem 2.6), if the *a*_*i*_ are fixed with *a*_1_ ⩾ *a*_2_ ⩾ … ⩾ *a*_*I*_ and the *b*_*i*_ are free to vary subject to *b*_1_ ⩾ *b*_2_ ⩾ … ⩾ *b*_*I*_ (and 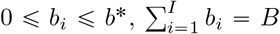), then for each permutation *σ* of (1, 2, …, *I*),

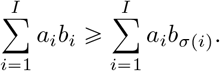

In other words, to maximize 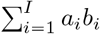, it suffices to proceed by assuming that *b*_1_ ⩾ *b*_2_ ⩾ … ⩾ *b*_*I*_.

We apply Lemma E.2 with *ℓ* = 1. We conclude that the maximal value of 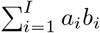 is achieved with *b*_1_ = min(*b*\*, *B*) = *b*\*. Fixing *b*_1_ = *b*\*, we next apply Lemma E.2 with *ℓ* = 2. We find that the maximal value of 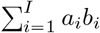 achieved at *b*_2_ = min(*b*\*, *B* − *b*\*).

We proceed by fixing *b*_*ℓ*_ = *b*\* for each *ℓ* = 3, 4, …, *⌈b*/*b*\* ⌉ − 1, repeatedly applying Lemma E.2 provided *ℓb*\* < *B*—that is, while *ℓ* < *B*/*b*\*, or *ℓ* ⩽ *⌈B*/*b*\* ⌉ − 1, and continuing to assign *b*_*ℓ*_ = *b*\*. The next value of *ℓ* is *ℓ* =. If ⌈*B*{*b*\*s ⩽ *I* − 1, then Lemma E.2 yields *b*_*ℓ*_ = *B* − *⌈B*/*b*\* ⌉ − 1 *b*\* and *b*_*i*_ = 0 for all *i* > *ℓ*. If *⌈B*/*b*\* ⌉ = *I*, then we have reached a trivial case in which *b*_*I*_ = *B* −(*I* − 1)*b*\* and *b*_*i*_ = *b*\* for all *i*, 1 ⩽ *i* ⩽ *I* − 1.

We arrive at Condition 3: with the *a*_*i*_ in non-increasing order held constant, the *b*_*i*_ that satisfy Condition 3 produce the maximum of 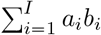. By symmetry, we fix the *b*_*i*_ as in Condition 3 and apply Lemma E.2 with the roles of the *a*_*i*_ and *b*_*i*_ interchanged. We find analogously that the *a*_*i*_ follow Condition 1.

(ii) For the lower bound, by the rearrangement inequality (Theorem 2.6), if the *a*_*i*_ are fixed with *a*_1_ ⩾ *a*_2_ ⩾ … ⩾ *a*_*I*_ and the *b*_*i*_ are free to vary subject to *b*_1_ ⩽ *b*_2_ ⩽ … ⩽ *b*_*I*_ (and 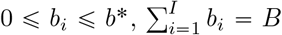), then for each permutation *σ* of (1, 2, …, *I*),

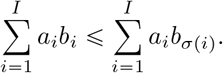

In other words, to minimize 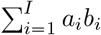, it suffices to proceed by assuming that *b*_1_ ⩽ *b*_2_ ⩽ … ⩽ *b*_*I*_.

We apply Lemma E.1 with *ℓ* = *I*. We conclude that the minimal value of 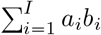 is achieved with *b*_*I*_ = min(*b*\*, *B*) = *b*\*. Fixing *b*_*I*_ = *b*\*, we next apply Lemma E.1 with *ℓ* = *I* − 1. We find that the minimal value of 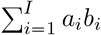 is achieved at *b*_*I*−1_ = min(*b*\*, *B* − *b*\*).

We proceed by fixing *b*_*ℓ*_ = *b*\* for each *ℓ* = *I* − 2, *I* − 3, …, *I* − ⌈*B*/*b*\* ⌉ + 2, repeatedly applying Lemma E.1 provided 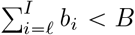, that is, while (*I* − *ℓ* + 1)*b*\* < *B*, or *ℓ* ⩾ *I* − ⌈*B*/*b*\* ⌉ + 2, and continuing to assign *b*_*ℓ*_ = *b*\*.

The next value of *ℓ* is *ℓ* = *I* − ⌈*B*/*b*\* ⌉ + 1. If *I* − ⌈*B*/*b*\* ⌉ + 1 ⩾ 2, then Lemma E.1 yields *b*_*ℓ*_ = *B* − (⌈*B*/*b*\* ⌉ − 1)*b*\*, and *b*_*i*_ = 0 for all *i* < *ℓ*. If *I* −⌈*B*/*b*\* ⌉ + 1 = 1, then we have reached a trivial case in which *b*_1_ = *B* −(*I* − 1)*b*\* and *b*_*i*_ = *b*\* for all *i*, 2 ⩽ *i* ⩽ *I*.

We arrive at Condition 2: with the *a*_*i*_ in non-increasing order held constant, the *b*_*i*_ that satisfy Condition 2 produce the minimum of 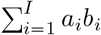. By symmetry, if we fix the *b*_*i*_ as in Condition 2, and write the *b*_*i*_ in reverse order, with 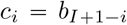, then we can apply Lemma E.1 with the *c*_*i*_ in the role of the *a*_*i*_ and the reversed *a*_*i*_, or 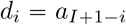 in the role of the *b*_*i*_. We obtain that the *d*_*i*_ follow Condition 2, and consequently, that the 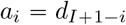 follow Condition 1.

**Proof of Theorem 4.5** The function 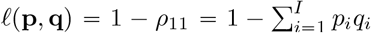, With 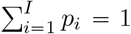 and 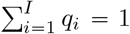, is minimized when 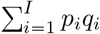 is maximized, and maximized when 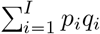 is minimized.

i. Via Lemma E.3i, 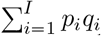 reaches its upper bound if 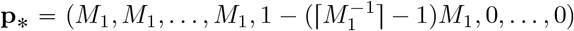 and 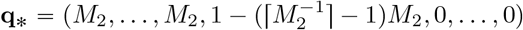, producing the lower bound for *ℓ*(**p, q**).
ii. Via Lemma E.3ii, 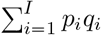 reaches its lower bound if 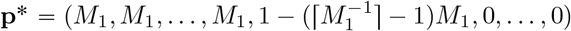 and 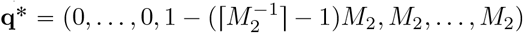, producing the upper bound for *ℓ*(**p, q**).

The values of *ℓ*(**p**_∗_, **q**_∗_) and *ℓ*(**p**\*, **q**\*) can be obtained by computing *ℓ*(**p, q**) with the vectors specified.

## F Proof of Corollary 4.6

This proof follows that of Theorem 4.5. With the additional requirement that *M*_1_ and *M*_2_ are the frequencies for the same allele in both populations, we can write *ℓ*(**p, q**) as,

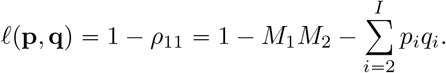

We must find (*p*_2_, *p*_3_, …, *p*_*I*_) and (*q*_2_, *q*_3_, …, *q*_*I*_) that give the upper and lower bounds for 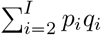.

With 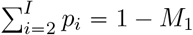 and 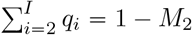, by Lemma E.3, the minimum of 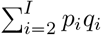 is reached at

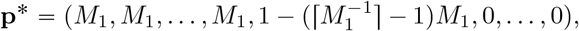

with 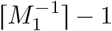 entries of *M*_1_ followed by an entry of 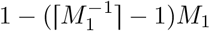 and 0 for the rest, and

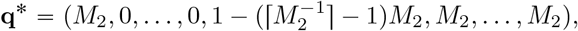

with one entry of 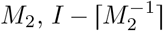 entries of 0, followed by an entry of 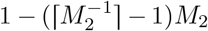 and 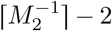 entries of *M*_2_. These values minimize 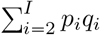, thereby maximizing 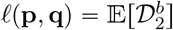

Similarly, by Lemma E.3, the maximum of 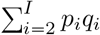 is reached at

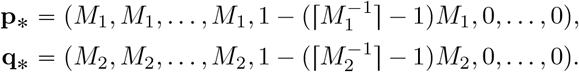

These values maximize 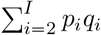, thus minimizing 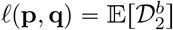.

The values of *ℓ*(**p**\*, **q**\*) and *ℓ*(**p**_∗_, **q**_∗_) can be obtained accordingly.

